# Inhibition of pyrimidine synthesis in murine skin wounds induces a pyoderma gangrenosum-like neutrophilic dermatosis accompanied by spontaneous gut inflammation

**DOI:** 10.1101/2022.12.20.521286

**Authors:** Samreen Jatana, András K. Ponti, Erin E. Johnson, Nancy A. Rebert, Jordyn L. Smith, Clifton G. Fulmer, Edward V. Maytin, Jean-Paul Achkar, Anthony P. Fernandez, Christine McDonald

## Abstract

Pyoderma gangrenosum (PG) is a debilitating skin condition often accompanied by inflammatory bowel disease (IBD). Strikingly, ∼40% of patients that present with PG have underlying IBD, suggesting shared but unknown pathogenesis mechanisms. Impeding the development of effective treatments for PG is the absence of an animal model that exhibits features of both skin and gut manifestations. This study describes the development of the first experimental drug-induced mouse model of PG with concurrent intestinal inflammation. Topical application of pyrimidine synthesis inhibitors on wounded mouse skin generates skin ulcers enriched in neutrophil extracellular traps (NETs) and pro-inflammatory cellular as well as soluble mediators mimicking human PG. The mice also develop spontaneous intestinal inflammation demonstrated by histologic damage. Further investigations revealed increased circulating immature low-density IL-1β primed granulocytes that undergo enhanced NETosis at inflamed tissue sites supported by increase in circulatory citrullinated histone 3, a marker of aberrant NET formation. Granulocyte depletion dampens the intestinal inflammation in this model, further supporting the notion that granulocytes contribute to the skin-gut crosstalk in PG mice. We anticipate that this novel murine PG model will enable researchers to probe common disease mechanisms and identify more effective targets for treatment for PG patients with IBD.

## Introduction

Pyoderma gangrenosum (PG) is a rare, sterile, neutrophilic dermatosis of the skin. PG is a common cutaneous manifestation of inflammatory bowel disease (IBD), more frequently associated with ulcerative colitis than Crohn’s disease (1-3). Approximately 30-40% of patients with PG have underlying IBD and a large proportion of them observe PG onset before their diagnosis of IBD is confirmed (4-7). Multiple syndromic forms of PG exist, but the most common ulcerative form of PG typically begins as an erythematous nodule that evolves into a chronic purulent ulcer with violaceous undermined borders (8-10). PG lesions are often preceded by trauma to the skin, a phenomenon known as pathergy (11). Common sites of PG lesions include the extremities, genitalia, perineal area, and postsurgical stoma sites (8, 12). While the course of other cutaneous manifestations associated with IBD, such as erythema nodosum and Sweet’s syndrome, often run parallel to that of IBD activity, PG can occur before IBD diagnosis or manifest even after IBD is in remission (4, 5, 13).

The factors driving PG pathogenesis and the underlying mechanisms that link it to IBD remain elusive. This is partly due to limited data available from human studies as well as the lack of animal models with concurrent skin and intestinal inflammation. Genome-wide association studies have identified a number of susceptibility loci conferring the risk of development of PG in patients with IBD, including loci that play a role in neutrophil recruitment (IL8RA), IL-17-mediated cellular immune responses (TRAF3IP2), and extracellular matrix degradation (TIMP3) (7, 14). Additional studies examining individuals with PG and its syndromic form PG, acne and suppurative hidradenitis (PASH) have identified mutations in components involved in regulation of innate immune responses (NLRP3, NLRP12, MEVF, NOD2, PSTPIP1), implicating dysregulated molecular pattern recognition signaling (15, 16). Of note, the hereditary autosomal dominant form of PG known as pyogenic arthritis, PG and acne (PAPA syndrome) is associated with mutations in proline–serine–threonine phosphatase interacting protein 1 (PSTPIP1) (17). Mutations in PSTPIP1 can cause pyrin-mediated activation of the inflammasome due to hyperphosphorylation of PSTPIP1 leading to increased production of pro-inflammatory factors like IL-1β and IL-18, both of which have been implicated in PG pathogenesis (18-20). PG patients often benefit from treatment with anakinra, an interleukin-1 receptor antagonist (IL-1Ra), further supporting the role of dysregulated innate immune responses (9). Mice ectopically expressing the PAPA-associated PSTPIP1 A230T mutant protein show elevated levels of circulating cytokines implicated in active PG, but they fail to develop skin inflammation and arthritis, specific to PAPA syndrome (21). Other insights into disease pathogenesis have been provided by animal models of neutrophilic dermatosis with mutations of tyrosine-protein phosphatase non-receptor type 6 gene (PTPN6), specifically highlighting the role of IL-1R signaling and apoptosis signal-regulating kinases in disease progression (22-24). While findings from both clinical and animal models have improved our understanding of PG pathobiology, there are major gaps in identifying the inflammatory triggers that contribute to skin-gut crosstalk in patients with PG and IBD (25). None of the existing genetically driven models have demonstrated the inter-organ communication between the skin and intestine.

In this study, we present a novel, drug-induced mouse model of PG-like neutrophilic dermatosis with concomitant intestinal inflammation. Our data show that the topical application of pyrimidine synthesis inhibitors to murine skin wounds generates non-healing skin ulcers enriched in neutrophils mimicking the PG disease phenotype. Unlike existing PG animal models, these mice also display spontaneous intestinal inflammation indicating the existence of a pathogenic inflammatory crosstalk between the skin and the gut. Our results demonstrate that priming and activation of immature, low-density granulocytes with IL-1β in diseased animals leads to exaggerated neutrophil extracellular trap (NET) formation, as shown by the presence of NETs at sites of inflammation in both the skin as well as the intestine. Importantly, we also demonstrate that depletion of granulocytes reduces intestinal inflammation suggesting that uncontrolled neutrophil activation and migration to sites of inflammation driven by chemokine cues is a key pathogenic mechanism in this novel PG model.

## Results

### Skin-wounded mice treated topically with a pyrimidine synthesis inhibitor exhibit a PG-like neutrophilic dermatosis

Pyrimidine and purine metabolism have been targeted in various diseases to elicit immunomodulatory responses (26-31). Clinical studies have shown that azathioprine, an inhibitor of purine synthesis, utilized for the treatment of IBD, can precipitate neutrophilic dermatosis such as Sweet syndrome and PG in skin (32-34). In this study, we utilized topical pyrimidine synthesis inhibition in mouse wounds to induce a neutrophilic dermatosis phenotype. Wild-type C57BL/6J with circular full-thickness excisional skin wounds were treated daily for a period of 9 days with topical 2% N-phosphonacetyl-L-aspartate (PALA) (**Figure 1A**). PALA is a specific transition state inhibitor of a trifunctional protein, carbamoyl phosphatase synthase II/aspartate transcarbamylase/dihydroorotase (CAD), which is the enzyme required for the first three steps of the pyrimidine synthesis pathway (**Figure 1B**). Mice treated with topical PALA developed non-healing purulent ulcers in the wounded region compared to control mice treated with Aquaphor alone (**Figure 1C**). These PALA-induced phenotypical changes were observed starting at day 4 and were fulminant by day 9. Detailed histological analysis of tissue sections revealed deleterious inflammatory changes including the complete loss of the epidermis and a large number of immune cell infiltrates in the ulcerated region of mice treated with PALA (**Figure 1D**). In contrast, wounds in mice treated with Aquaphor alone had completely healed by contraction as evidenced by the presence of a distinct hyperproliferative epidermal layer and obvious granulation tissue in the wound area by day 9 (**Figure 1D**).

**Figure 1.**
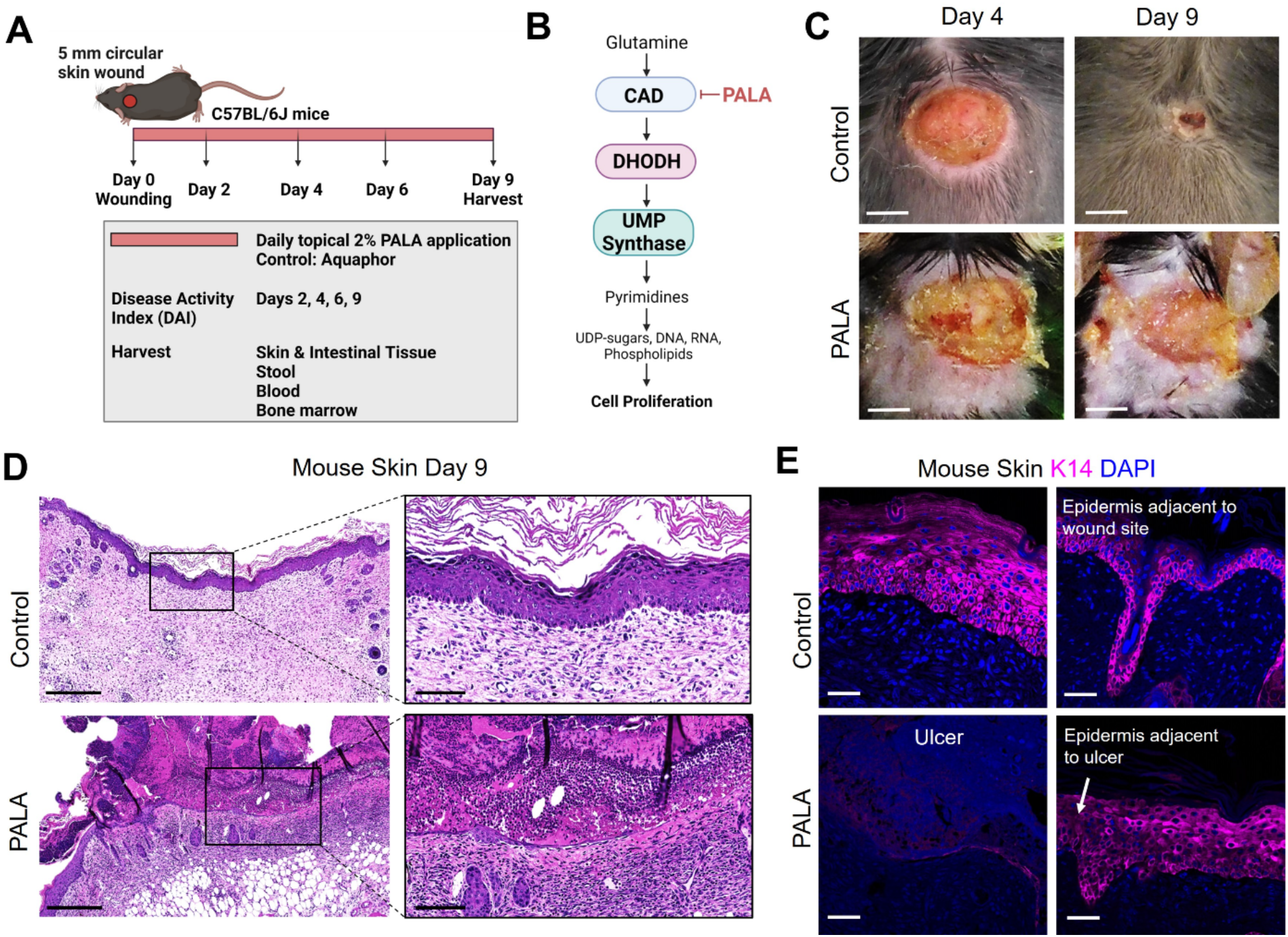
Skin-wounded mice treated topically with a pyrimidine synthesis inhibitor exhibit a PG-like neutrophilic dermatosis. (**A**) *In vivo* experimental set-up and treatment timeline in the PG mouse model. (**B**) Mechanism of pyrimidine synthesis inhibition by N-phosphonacetyl-L-aspartate (PALA). (**C**) Visual appearance of PG-like ulcers in mouse skin photographed on day 4 and day 9 post-wounding. Scale bars: 0.2 cm. (**D**) Histopathology of skin tissue sections stained with H&E on day 9 post-wounding. Scale bars: 500μm, inset: 100μm. (**E**) Immunofluorescence (IF) staining of keratin 14 (K14, magenta) in mouse skin in the wound region (left) and adjacent to the wound site (right). Tissue harvested on day 9 post-wounding. Scale bars: 50μm. Nuclei are stained with DAPI (blue).

Human PG ulcers show a characteristic undermined border and dense neutrophilic infiltrates that damage both the epidermis and dermis as the ulcer expands due to inflammation (35). In order to specifically visualize the epidermal layer, immunofluorescence staining was performed on murine skin tissue to identify keratin 14 (K14), a cytoplasmic intermediate filament protein expressed in mitotically active basal keratinocytes of the epidermis (36, 37). K14 staining showed an absence of the epidermal layer in the ulcer region of the PALA-treated mice, as compared to control mice that had high skin K14 expression in the proliferative epidermal region of the wound site (**Figure 1E**). The epidermal region adjacent to the wound site in PALA-treated mice showed expression of K14 in proliferative keratinocytes (**Figure 1E**). In this model, application of topical PALA inhibited the epidermal cell regeneration and migration, which is an essential step in the wound healing process. Overall, there is conspicuous histopathological similarity between the ulcers observed in the wounded mouse skin treated with PALA and the clinical histopathology of human PG.

### Similar cellular and soluble mediator inflammatory landscape is observed in murine PG and human PG

The inflammatory landscape in human PG has been well characterized (9). In human PG, dense neutrophilic infiltrates are typically found in the ulcer and underlying dermis, accompanied by perivascular monocytes and lymphocytes in the region peripheral to the ulcer (9). In order to qualitatively evaluate the immune cell infiltrates in PALA-treated mouse ulcers, skin tissue was probed using immunofluorescence staining with MPO, F4/80 and CD3 to visualize neutrophils, macrophages and T cells, respectively. PALA-treated skin revealed the presence of abundant neutrophils in the ulcer region and underlying dermis, compatible with the role of PALA in inducing a murine neutrophilic dermatosis closely resembling human PG (**Figure 2A**). Recurrent topical PALA application over 9 days arrested the wound healing process in the initial inflammatory phase leading to massive neutrophilic inflammation that resulted in the formation of highly exudative non-healing skin ulcers. In contrast, an increased number of macrophages were present in the skin of control mice, indicating that, in the absence of PALA, the wounds had progressed as in a normal wound healing process (**Figure 2A**). An overview of the PALA-treated tissue revealed that while neutrophils were predominant in the ulcerated region, dermis and regions surrounding the ulcer, macrophages were present in the deeper region of the dermis and the periphery of the wound (**Figure 2B**). Taken together, these results show that topical application of a pyrimidine synthesis inhibitor on murine skin wounds leads to robust cutaneous neutrophilic inflammation and stalling of the wound healing process, which accurately mimics the histopathology of human PG based on gross and histological evaluation.

**Figure 2.**
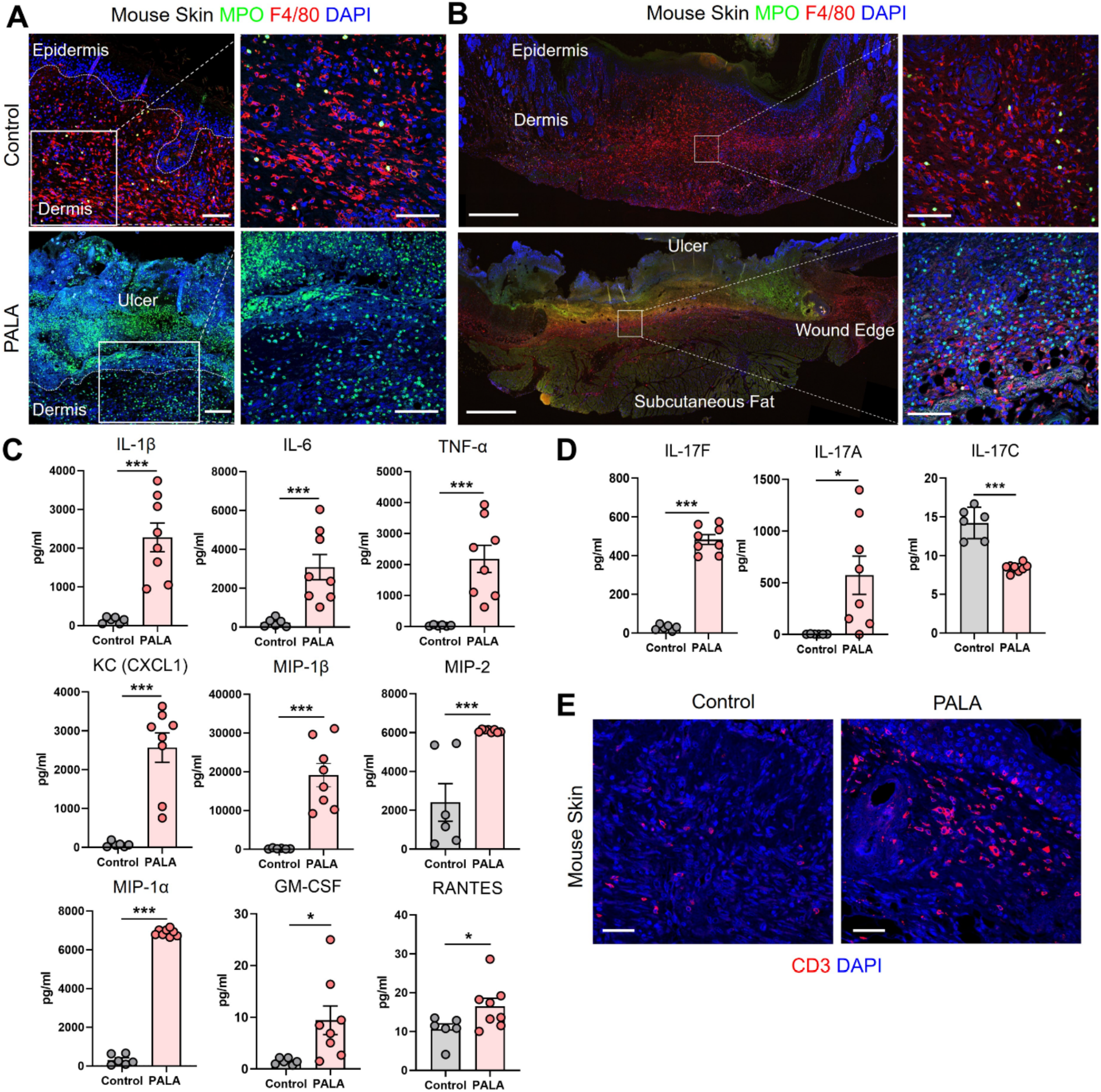
Similar cellular and soluble mediator inflammatory landscape is observed in murine PG and human PG. (**A**) IF staining of MPO (green) for neutrophils and F4/80 (red) for macrophages in the wounded region of the skin in mice treated with Aquaphor (control) and PALA. Scale bars: 50μm, inset: 50μm. (**B**) IF visualization of the entire cross-section of the skin tissue stained for neutrophils (MPO, green) and macrophages (F4/80, red). Scale bars: 500μm, inset: 50μm. (**C**) Quantification of the cytokines (IL-1β, IL-6, TNF-α) and chemokines (CXCL1, MIP-1β, MIP-2, MIP-1α, GM-CSF, RANTES) in skin tissue homogenates. (**D**) Quantification of Th17 cytokines (IL-17F, IL-17A, IL-17C) in skin tissue homogenates. (**E**) IF staining of CD3+ T cells (red) in mouse skin tissue. Scale bars: 50μm. Nuclei in all IF images are stained with DAPI (blue). All data is presented as Mean ± SEM, n=8, statistical significance determined by unpaired, nonparametric, two-tailed Mann Whitney test. *p<0.05, ***p<0.001.

Given the striking similarities of the cellular infiltrates of murine PG lesions with human ones, we next explored whether a similar pattern of soluble inflammatory mediators was also present. In order to characterize the expression profile of soluble inflammatory mediators in the wounded mouse skin, skin tissue homogenates were analyzed by multiplexed cytokine and chemokine analysis. The analytes selected for the multiplex panel were based on the well documented inflammatory milieu contributing to disease pathogenesis in human PG (9, 15, 38). The expression of pro-inflammatory cytokines including interleukin 1L-1β (IL-1β), IL-6, and tumor necrosis factor-α (TNF-α) was significantly higher in PALA-treated mice compared to controls (**Figure 2C**). The levels of neutrophil, macrophage and T cell chemoattractants including chemokine (C-X-C motif) ligand 1 (CXCL1), macrophage inflammatory protein-2 (MIP-2), MIP-1α, MIP-1β, granulocyte-macrophage colony stimulating factor (GM-CSF) and RANTES (CCL5) were also significantly higher in PALA-treated mice compared to controls (**Figure 2C**).

Lastly, elevated levels of IL-17A and IL-17F were also found in skin tissue of PALA-treated mice (**Figure 2D**). Interestingly, IL-17C, which is mainly produced by epithelial cells, was significantly downregulated on day 9 probably due to the loss of the epidermal layer in PALA-treated mice (**Figure 2D**). To complement these observations of enriched inflammatory mediators, we performed immunofluorescence staining to qualitatively evaluate the presence of T cells in mouse skin tissue and detected abundant CD3+ T cells in the vicinity of the ulcer region in the dermis of PALA-treated mice (**Figure 2E**). These results demonstrate biological similarities in the inflammatory milieu between the PG phenotype in mice and human disease (9). The inflammatory mediators we detected produced by epidermal, dermal, and immune cells, suggests that both the innate and adaptive immune responses drive the PALA-induced inflammatory cascade in our murine PG model.

### Mice with PG phenotype develop spontaneous intestinal inflammation dependent on the presence of skin wound

There is significant epidemiological overlap in the concurrent development of PG and IBD (39), raising the key question of their pathophysiological interdependence. During the 9-day treatment regimen, disease activity index (DAI) was measured in the mice. DAI assessment comprised of change in body weight, posture (normal vs. hunched), fur (normal vs. ruffled), stool consistency and evaluation of rectal prolapse every 2-3 days (**Supplemental Table 1**). While PALA-treated mice did not experience significant weight loss, their DAI scores increased over time and were significantly higher in comparison to control mice on day 9 (**Figure 3A**). Noticeable changes to the stool consistency were observed during this period. As a result, the colon and terminal part of the ileum were harvested from mice on day 9 and histopathological assessment was performed to evaluate the presence of intestinal injury that included key features such as presence of inflammatory infiltrates, neutrophils, crypt density, crypt hyperplasia, goblet cell loss, submucosal swelling, muscle layer thickening, presence of crypt abscess and ulceration (**Supplemental Table 2**) (40). In PALA treated mice, a range of inflammation was observed in the distal colon indicated by inflammation scores that extended from 6 to 24 (**Figure 3B and 3D, Supplemental Figure 1A**). The most severe inflammation observed in the colon was characterized by the loss of crypt structures, ulceration of the epithelial cell layer evident by E-cadherin staining, and pronounced immune cell infiltrates (**Figure 3B and 3C**). Inflammation scores were significantly higher in the transverse and distal colon of PALA-treated mice compared to controls, suggesting that inflammation was more prominent in the lower half of the colon (**Figure 3D**). Significant from a temporal perspective, development of intestinal lesions trailed behind skin inflammation in our model, as colonic tissue collected on day 4 displayed minimal inflammation compared to controls but the skin inflammation was already present at day 4 (**Supplemental Figure 1B and 1C**).

**Figure 3.**
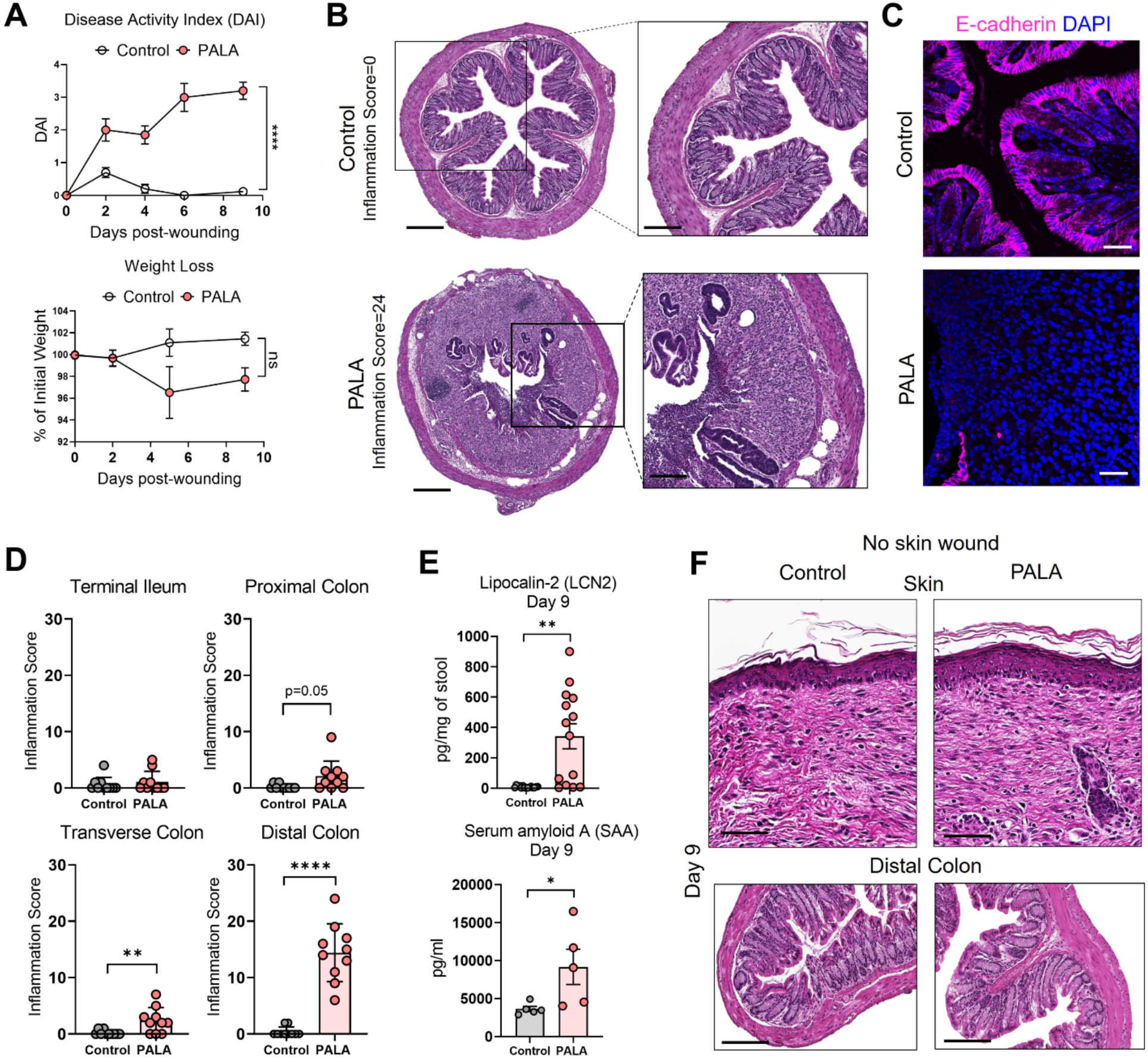
Mice with PG phenotype develop spontaneous intestinal inflammation dependent on the presence of skin wound. (**A**) Disease activity index (DAI) and weight loss assessment in mice over a period of 9 days. (**B**) Histopathology of the cross section of the distal colon stained with H&E on day 9 post-wounding. Scale bars: 300μm, inset: 100μm. (**C**) E-cadherin staining (magenta) in distal colon tissue. Nuclei are stained with DAPI (blue). Scale bars: 50μm. (**D**) Assessment of the inflammation score in mice treated with PALA compared to controls in the terminal ileum, proximal colon, transverse colon and distal colon. (**E**) Fecal Lcn-2 and serum SAA quantification utilizing ELISA in the stool and serum of mice, respectively. (**F**) Histopathology of skin (top) and distal colon (bottom) of mice treated with topical PALA and Aquaphor (control) without the presence of a skin wound for a duration of 9 days. Scale bars: 100μm. Data is presented as Mean ± SEM, n=10-25 (**A**), n=5-14 (**D, E**), statistical significance determined by unpaired, nonparametric, two-tailed Mann Whitney test. *p<0.05, **p<0.01 and ****p<0.0001.

Additional parameters of local and systemic inflammation were also measured, including fecal lipocalin-2 (Lcn-2) and serum amyloid A (SAA) in stool and blood, respectively. Fecal Lcn-2 is a non-invasive biomarker used to detect intestinal inflammation in mice, similar to fecal calprotectin utilized to evaluate intestinal inflammation in human stool, while SAA, is an acute phase protein synthesized by the liver (41, 42). Fecal Lcn-2 levels were significantly increased in mice treated with PALA, further corroborating the results from the gut histopathology analysis (**Figure 3E**). SAA levels were significantly increased in PALA-treated mice compared to controls demonstrating that in addition to local tissue damage in the skin and gut, these mice also had elevated systemic inflammation, as commonly seen in patients with IBD (**Figure 3E**).

Skin insult or pathergy is a common event in individuals that develop PG (8). To determine whether a skin insult is required for the development of murine PG and concomitant intestinal inflammation, topical PALA was applied to intact mouse skin for 9 days. Topical PALA application in the absence of a skin insult failed to generate a local inflammatory response in the skin and development of concomitant colonic inflammation (**Figure 3F**). In addition, no differences were observed in the fecal Lcn-2, systemic SAA and IL-1β (skin and colon) levels in these animals (**Supplemental Figure 1D**). These results indicate that persistent inflammation in the skin of mouse wounds treated with PALA precedes and seemingly leads to the development of intestinal inflammation mainly localized to the colon. The results demonstrate that both the presence of a skin insult and inhibition of pyrimidine synthesis by PALA is required for the development of PG-like inflammation in mouse skin and concomitant colonic inflammation.

### Inflammatory milieu in the distal colon of PG mice resembles UC and reflects systemic inflammation

Next, the cellular inflammatory environment in the distal colon of PG mice was assessed. Visualization and qualitative assessment of neutrophils (MPO), macrophages (F4/80) and T cells (CD3) using immunofluorescence staining showed an increase in these specific cell populations in the distal colon of PG mice compared to control (**Figure 4A**). To better characterize the soluble mediators of the inflammatory landscape of the intestine, cytokines and chemokines were quantified in tissue homogenates of the distal colon. Pro-inflammatory cytokines IL-1β, TNF-α and IL-6 were significantly elevated in PALA-treated mice compared to controls (**Figure 4B**). Colonic levels of chemoattractants including CXCL1 (KC), monocyte chemoattractant protein-1 (MCP-1), MIP-1α, MIP-2 and GM-CSF were also significantly increased in PALA-treated mice (**Figure 4B**).

**Figure 4.**
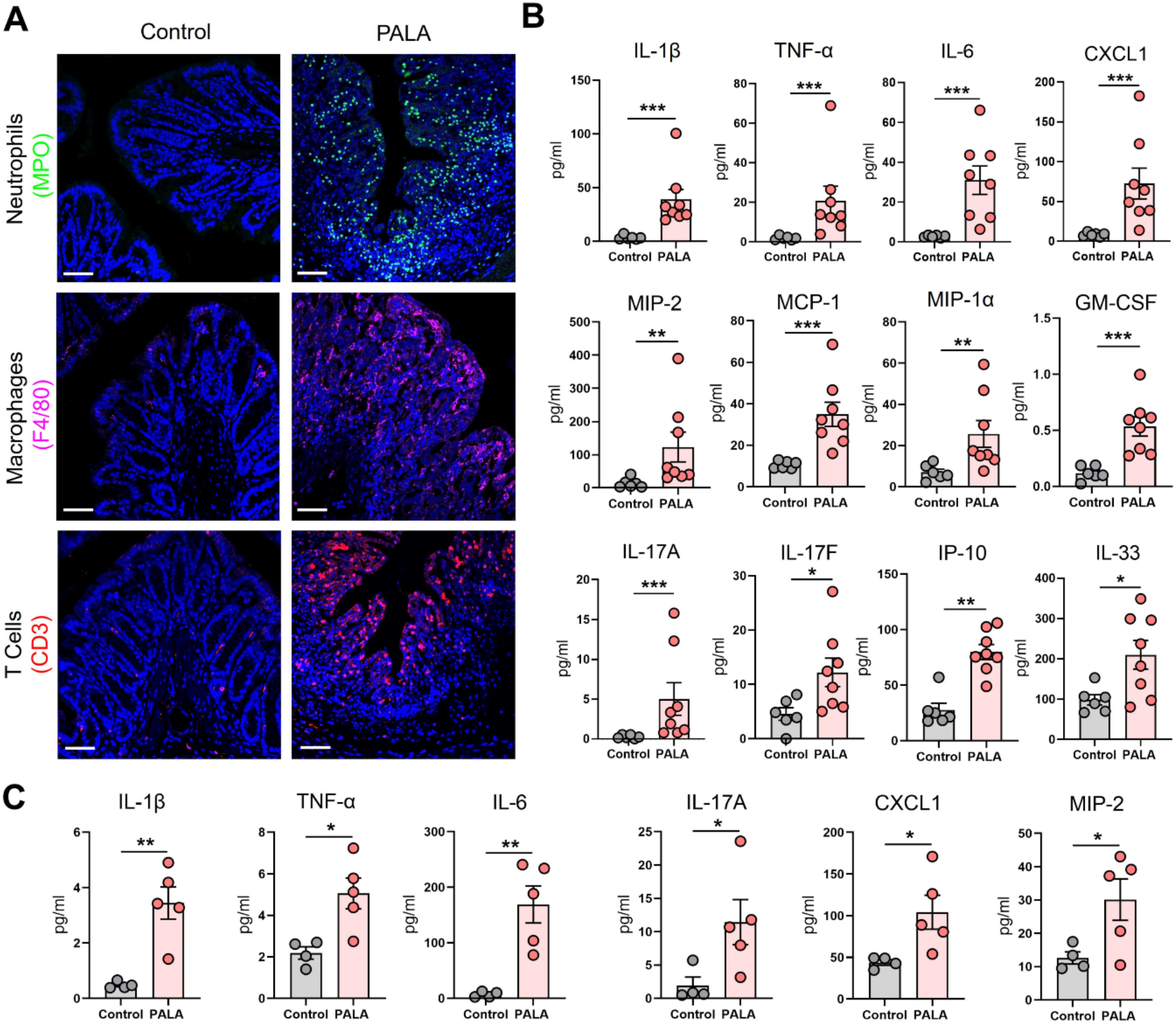
Inflammatory milieu in the distal colon of PG mice resembles UC and reflects systemic inflammation. (**A**) Qualitative assessment of presence of immune cell infiltrates including neutrophils (MPO, green), macrophages (F4/80, magenta) and CD3+ T cells (red) in the distal colon of PALA-treated mice compared to controls. Nuclei are stained with DAPI (blue). Scale bars: 50μm. (**B**) Quantification of cytokines and chemokines in tissue homogenates from the distal colon. (**C**) Quantification of cytokines and chemokines in the plasma of mice treated with PALA compared to control treatment. All data is presented as Mean ± SEM, n=4-8, statistical significance determined by unpaired, nonparametric, two-tailed Mann Whitney test. *p<0.05, **p<0.01, ***p<0.001.

IL-17C (epithelial) and IL-17A/F (T cells) cytokines were also elevated in the colons of PALA-treated mice, suggesting a Th17 phenotype in the distal colon similar to that observed in the PG mouse skin (**Figure 4B**). Interferon-γ-inducible protein-10 (IP-10 or CXCL10), which has been shown to play a role in inflammatory cell migration to the gut in ulcerative colitis (UC) (43-45), was significantly increased in the PALA-treated group (**Figure 4B**). IL-33, which is released in a full-length form upon tissue damage or injury to the gut epithelium and cleaved to form mature IL-33 by proteases released from neutrophils (46-48), was significantly elevated in PALA mice (**Figure 4B**).

Finally, systemic cytokines and chemokines found to be significantly elevated in the plasma of PALA-treated mice included IL-1β, TNF-α, IL-6, IL-17A, CXCL1, and MIP-2 (**Figure 4C**). The above results indicate that certain cytokines and chemokines elevated in both the skin and intestine are elevated in the systemic circulation and are likely mediating local as well as systemic inflammatory effects. In particular, chemotactic cytokines involved in the migration of neutrophils were significantly increased. This finding, combined with the massive neutrophilic infiltration noted in both the skin and colon of PG mice strongly suggests that neutrophils may be the key drivers of the inflammatory response observed in our PG model.

### IL-1β-driven priming of low-density granulocytes leads to NET formation and contributes to concomitant skin and intestinal inflammation

Neutrophils undergo a specialized form of cell death called neutrophil extracellular trap (NET) formation in inflammatory conditions like PG (49). Low-density granulocytes (LDGs), a distinct class of immature neutrophils, are elevated in the blood of individuals with PG and undergo spontaneous NET formation *in vitro* upon IL-1β priming (49). Therefore, the specific contribution of LDGs and NET formation in our murine PG model was investigated. As shown above, topical PALA application to wounded skin increased the influx of neutrophils to the skin and colon, but to determine how neutrophils specifically contribute to inflammation in our model, the presence of NETs in murine tissue was qualitatively assessed. Increased numbers of cells positive for citrullinated histone 3 (CitH3), a marker of NET formation (50), was observed in the ulcerated area of the skin and the inflamed distal colon of mice treated with PALA (**Figure 5A**).

**Figure 5.**
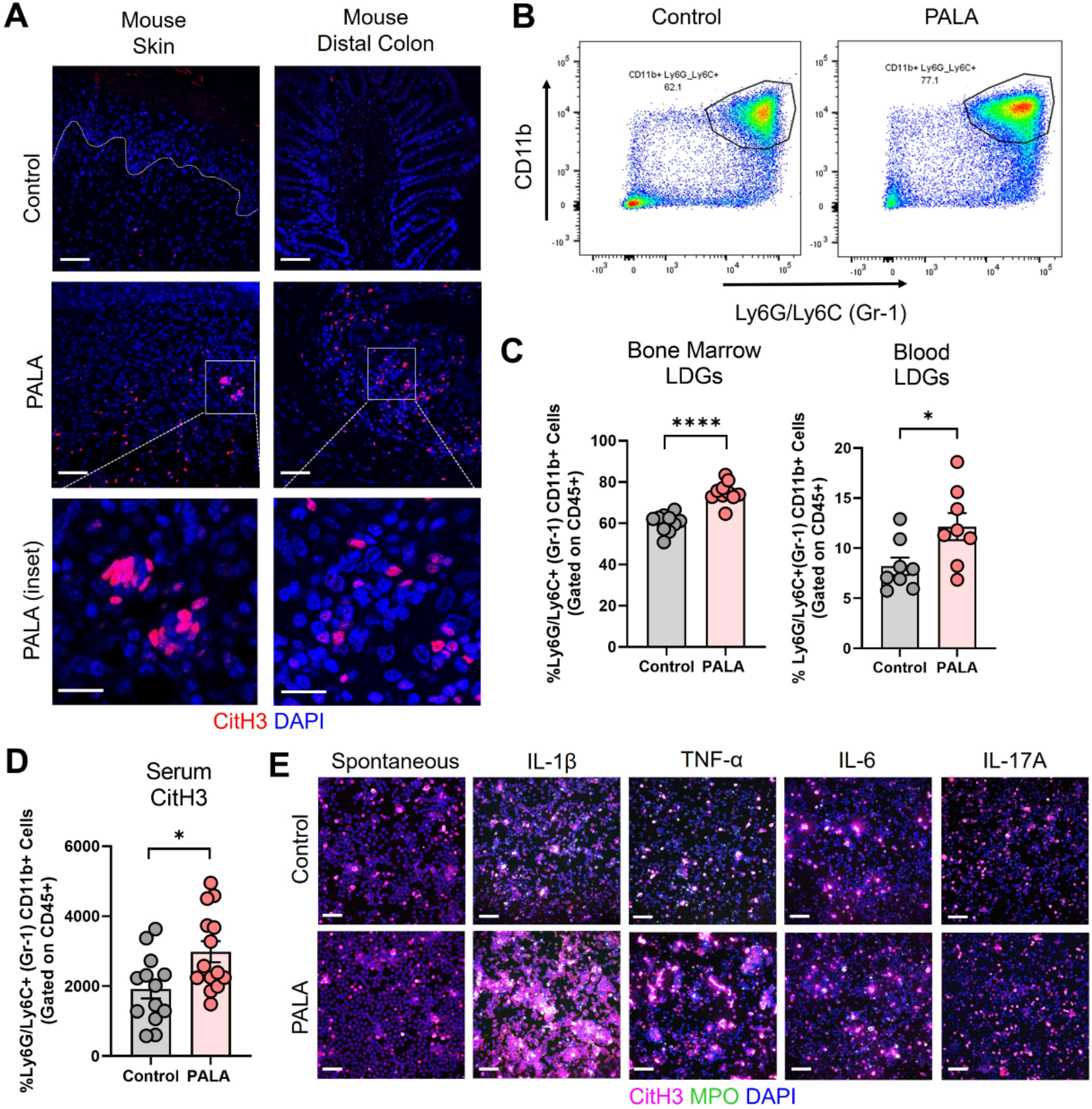
IL-1β-driven priming of low-density granulocytes leads to NET formation and contributes to concomitant skin and intestinal inflammation. (**A**) IF staining of CitH3 (red), a marker of NETs, in the skin and distal colon of mice treated with PALA compared to the control group. Nuclei are stained with DAPI (blue). Scale bars: 50μm, inset: 20μm. (**B**) Representative gating used to quantify LDGs in the blood and bone marrow of mice isolated using density gradient centrifugation. CD11b+ Gr-1+ cells were gated on CD45+ cells in the PBMC layer. (**C**) Quantification of LDGs in the bone marrow and blood of mice treated with PALA in comparison to the control group using flow cytometry. (**D**) Quantification of serum CitH3 levels in mice using ELISA. (**E**) IF staining and confocal imaging to quantify *in vitro* NET formation in LDGs isolated from the bone marrow. Groups include no treatment (spontaneous NETosis) and LDGs treated with IL-1β, TNF-α, IL-6 as well as IL-17A recombinant protein. NETs were stained with CitH3 (magenta), myeloperoxidase (MPO, green) and DAPI (blue). Scale bars: 50μm. All data is presented as Mean ± SEM, n=8-14, statistical significance determined by unpaired, nonparametric, two-tailed Mann Whitney test. *p<0.05 and ****p<0.0001. *In vitro* NETosis assays were performed in triplicate (n=3 animals), representative images shown in (**E**).

In order to further elucidate the role of neutrophils in the model pathobiology, we specifically investigated whether LDG or immature granulocyte production was enhanced since this subset of granulocytes has been shown to undergo exaggerated NETosis in PG (49). LDGs were isolated from the bone marrow and blood of the mice using density gradient centrifugation from the top layer of the gradient along with PBMCs and analyzed by flow cytometry (**Supplemental Figure 2A**). CD11b+ and Gr-1+ cells gated on CD45+ cells in the top gradient layer were quantified as the LDG population (**Figure 5B and Supplemental Figure 2B**). Both bone marrow and blood LDGs were significantly increased in PALA-treated mice, indicating that skin inflammation stimulates emergency granulopoeisis resulting in increased numbers of LDGs (**Figure 5C**). Moreover, serum CitH3 levels were significantly higher in PALA-treated mice compared to controls (**Figure 5D**), suggesting that immature neutrophils circulate and migrate based on chemotactic cues to distal sites where they undergo exaggerated NET formation that contributes to increased systemic CitH3 levels.

LDGs isolated from bone marrow were cultured *in vitro* to evaluate whether these cells are hyperactive and primed to undergo spontaneous NETosis or if a second pro-inflammatory hit is required for neutrophil activation at the tissue site. Pro-inflammatory mediators were selected from the systemic inflammatory profile of PALA-treated mice including IL-1β, TNF-α, IL-6 and IL-17A (**Figure 4C**). LDGs from PALA-treated mice did not undergo spontaneous NET formation; however, they did so in response to IL-1β stimulation *in vitro* (**Figure 5E**). Thus, LDGs in the bone marrow are primed to undergo exaggerated NET formation in the skin and distal colon, where elevated levels of IL-1β serve as a second hit to further induce NET formation. These results suggest that NET formation by activated LDGs is implicated in the PG-like inflammation and concomitant intestinal pathology in our model.

### Inhibition of pyrimidine synthesis is required for the induction of the murine PG-like phenotype and IL-1β-dependent NET formation

To determine whether the PALA-induced inflammatory phenotype in the skin is specifically a result of pyrimidine synthesis inhibition, another pyrimidine synthesis inhibitor was tested in this model. Brequinar is a quinolone carboxylic acid derivative that acts on dihydroorotate dehydrogenase (DHODH) to inhibit *de novo* pyrimidine synthesis (**Figure 6A**) (51). Topical brequinar was applied daily to excisional skin wounds to evaluate the inflammatory effects of pyrimidine synthesis inhibition on the skin and intestine. Similar to PALA-treated mice, topical brequinar (0.5% w/w)-treated animals also developed purulent, non-healing skin ulcers by day 9 (**Figure 6B and 6C**). Skin ulcers of brequinar-treated mice were enriched with neutrophils and peripheral macrophage infiltration (**Figure 6D**). In addition, increased CitH3 staining was observed indicating that neutrophils undergo enhanced NETosis at the ulcer site in response to topical brequinar (**Figure 6D**). These findings demonstrate that inhibition of pyrimidine synthesis in skin wounds drives the development of murine PG phenotype resembling human disease.

**Figure 6.**
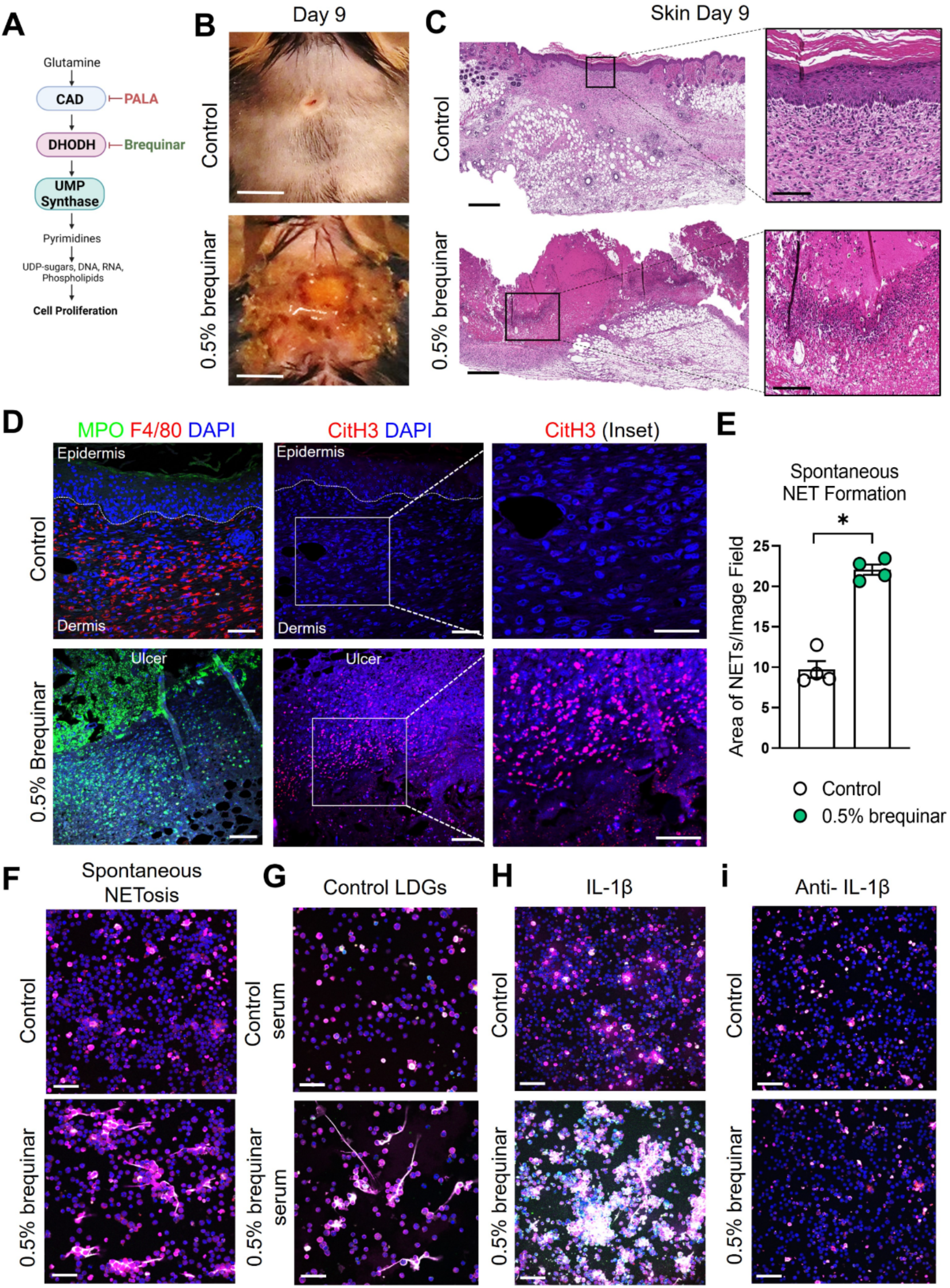
Inhibition of pyrimidine synthesis is required for the induction of the murine PG-like phenotype and IL-1β-dependent NET formation. (**A**) Mechanism of *de novo* pyrimidine synthesis inhibition of DHODH by brequinar. (**B**) Visual appearance of skin wounds in mice treated with 0.5% topical dose of brequinar compared to Aquaphor-treated controls on day 9 post-wounding. Scale bars: 0.5 cm. (**C**) Histopathology evaluation of mouse skin tissue stained with H&E on day 9 post-wounding. Scale bars: 500μm, inset: 100μm. (**D**) Qualitative assessment of neutrophils (MPO, green), macrophages (F4/80, red), NETs (CitH3, red) and DAPI (blue) in the wounded region of the skin treated with 0.5% brequinar compared to the control group. Scale bars: 50μm, inset: 50μm. (**E**) Quantification of *in vitro* spontaneous NET formation in LDGs isolated from the bone marrow of 0.5% brequinar-treated mice compared to the control group. Data is presented as Mean ± SEM, n=4, statistical significance determined by unpaired, nonparametric, two-tailed Mann Whitney test. *p<0.05. (**F**) Visualization of *in vitro* spontaneous NET formation in LDGs isolated from the bone marrow of 0.5% brequinar-treated mice. (**G**) *In vitro* NET formation was assessed in LDGs isolated from the bone marrow of control mice treated with control serum (top) and serum from 0.5% brequinar-treated mice (bottom). (**H**) IL-1β-induced NET formation in LDGs isolated from the bone marrow of 0.5% brequinar-treated mice in comparison to control. (**i**) Assessment of *in vitro* NET formation in the presence of neutralizing IL-1β antibody. NETs were stained with CitH3 (magenta), MPO (green) and DAPI (blue). *In vitro* NETosis assays were performed in triplicate (n=3 animals), representative images shown in (F-i), Scale bars (F-i): 50μm.

Interestingly, unlike PALA treatment, LDGs isolated from the bone marrow of brequinar-treated mice underwent spontaneous NET formation (**Figure 6E and 6F**). Enhanced NETosis was also observed in LDGs from control mice treated *ex vivo* with serum obtained from diseased brequinar-treated mice (**Figure 6G**). This result suggested that certain pro-inflammatory factors in systemic circulation contribute to enhanced priming of LDGs before they migrate to the inflamed tissue site, similar to the observation in PALA-treated mice. To identify pro-inflammatory factors involved in neutrophil priming and activation, LDGs isolated from the bone marrow of mice were treated with recombinant IL-1β, TNF-α, or IL-6. Treatment of LDGs with IL-1β exacerbated NETosis, leading to the formation of aggregated NETs *in vitro* suggesting that LDGs in brequinar-treated mice are primed with IL-1β (**Figure 6H and Supplemental Figure 3A**). Neutralization of IL-1β with anti-IL-1β antibody blocks this inflammatory process (**Figure 6i**). These results showed that LDGs are activated to undergo spontaneous NETosis and the addition of a second pro-inflammatory insult (IL-1β) exacerbates this phenomenon in brequinar-treated animals. These results establish that topical pyrimidine synthesis inhibition on wounded murine skin halts the wound healing process and is critical to generate murine neutrophilic dermatosis resembling human PG. NET formation in primed LDGs is further exacerbated by IL-1β, which contributes to chronic inflammation in the skin in this model.

### Intestinal inflammation induced by inhibition of pyrimidine synthesis depends on the type of inhibitor

Given the PG induction properties of brequinar, we investigated whether this type of pyrimidine synthesis inhibitor also induced intestinal inflammation. During the topical treatment period, DAI was measured in the animals treated with brequinar. Mice treated with topical 0.5% brequinar continued to lose body weight during the treatment period with significant changes at day 9 in comparison to controls (**Figure 7A**). DAI was also significantly higher in 0.5% brequinar-treated mice on harvest day (**Figure 7A**). Intestinal tissue from brequinar-treated mice was harvested to evaluate whether intestinal inflammation also developed as seen in PALA-treated animals. Histological analysis revealed that treatment of mouse wounds with topical 0.5% brequinar led to the development of low-level inflammation localized to the ileum characterized by presence of immune cells, decrease in villi length and submucosal swelling (**Figure 7B**). Thus, treatment of skin wounds with topical brequinar can also lead to the development of spontaneous intestinal inflammation, but in a different segment of the intestine, i.e., the ileum, in contrast to what was observed in PALA-treated mice, where intestinal inflammation was primarily evident in distal colon.

**Figure 7.**
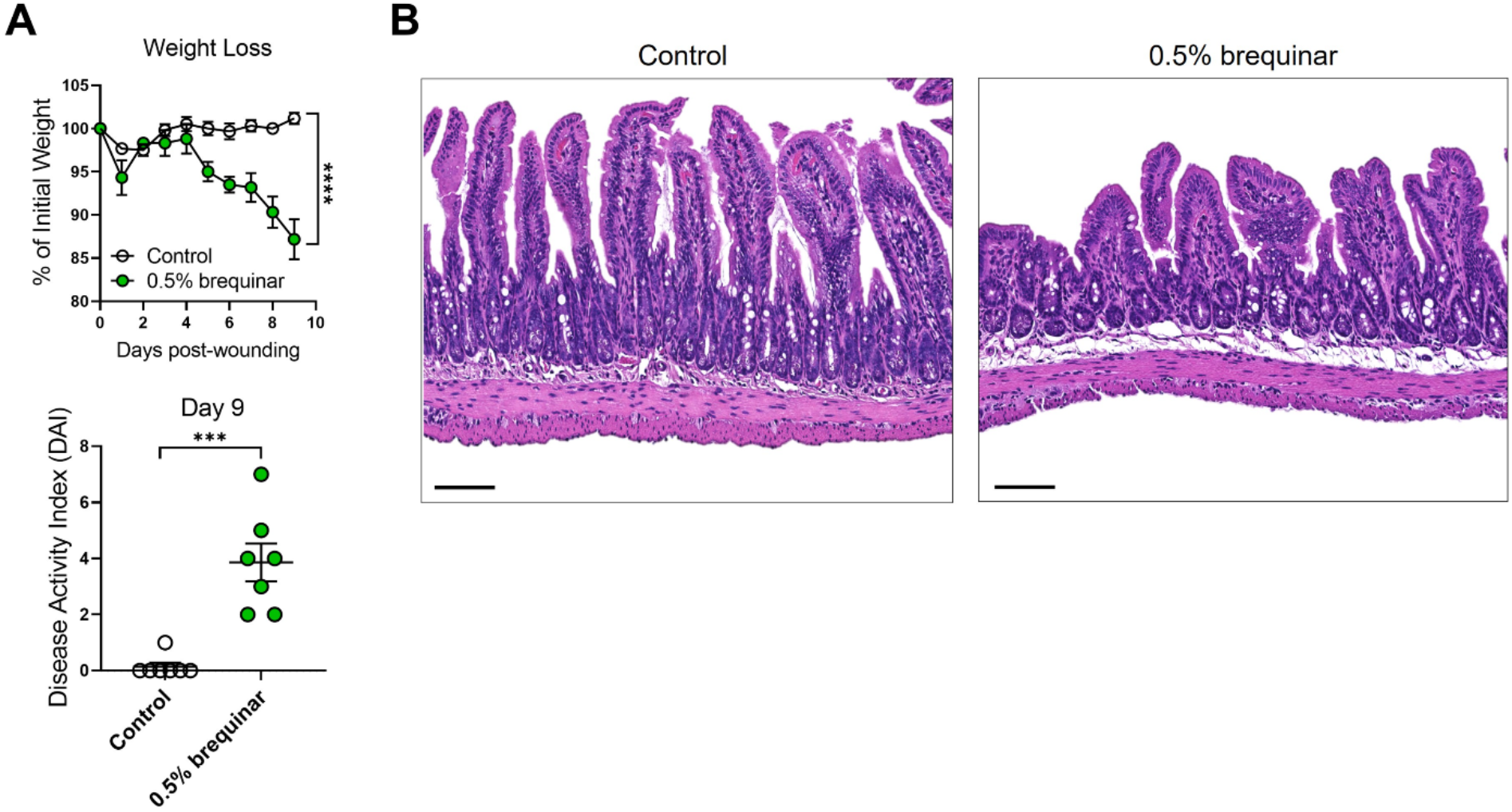
Intestinal inflammation induced by inhibition of pyrimidine synthesis depends on the type of inhibitor. (**A**) Weight loss and DAI assessment in mice treated with topical 0.5% brequinar over a period of 9 days compared to controls. (**B**) Histopathology of the cross section of the terminal ileum stained with H&E on day 9 post-wounding in topical 0.5% brequinar-treated animals. Scale bars: 100μm. All data is presented as Mean ± SEM, n=6-7, statistical significance determined by unpaired, nonparametric, two-tailed Mann Whitney test. ***p<0.001, ****p<0.0001.

### Granulocyte depletion mitigates intestinal inflammation in murine PG

The results of the NETosis assays performed in PALA- or brequinar-treated mice indicate that LDGs apparently play a key role in the pathogenesis of our novel model of murine PG. Therefore, the requirement of granulocytes in skin-gut crosstalk in our disease model was evaluated by depletion of granulocytes using anti-mouse Ly6G/Ly6C (Gr-1) antibody (**Supplemental Figure 4A**). Due to a high rate of mortality during granulocyte depletion in our longer 9-day treatment models (2% PALA and 0.5% brequinar), higher dose of topical brequinar (2% w/w) was utilized to induce significant skin and intestinal pathology rapidly over 6 days and reduce the number of animals required to achieve significant sample sizes. Mice treated with 2% topical brequinar had significant skin damage, underwent rapid weight loss, and had elevated DAI measurements by day 6 (**Figure 8, A-C**). Topical 2% brequinar-treated mice developed significant inflammation in the ileum by day 6 compared to mice treated with the lower dose 0.5% brequinar as evident by features like the presence of inflammatory cells, loss of epithelial layer, loss of goblet cells, decreased crypt density and shortened villi length (**Figure 8D and 8E**).

**Figure 8.**
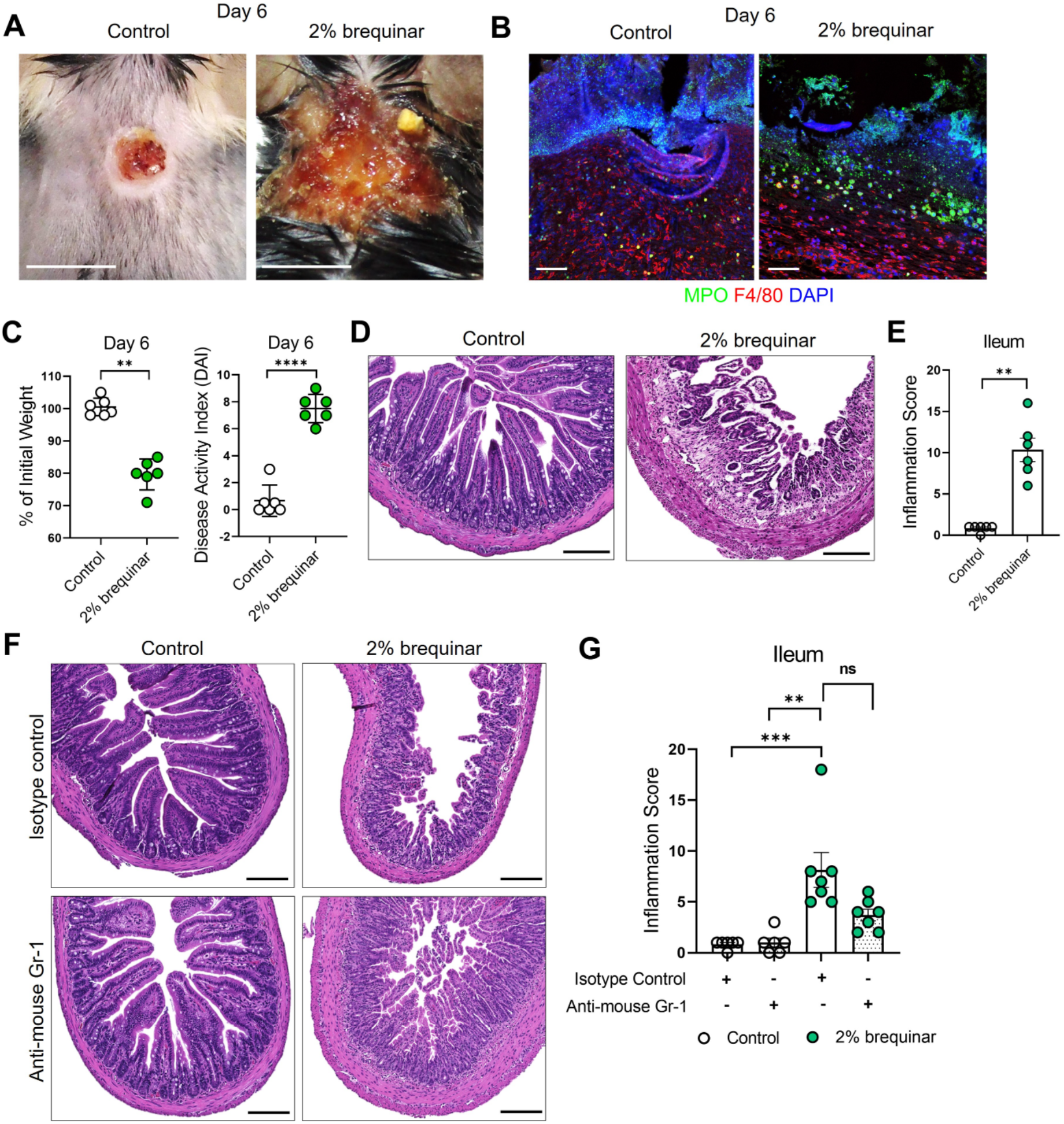
Granulocyte depletion mitigates intestinal inflammation in murine PG. (**A**) Visual appearance of wounds in the skin of topical 2% brequinar-treated mice compared to controls, 6 days post-wounding. Scale bars: 0.5 cm. (**B**) Qualitative assessment of the presence of neutrophils (MPO, green) and macrophages (F4/80, red) in the ulcer region of 2% brequinar-treated mice in comparison to control. Scale bars: 50μm. (**C**) Weight loss and disease activity index (DAI) assessment in mice treated with 2% brequinar over a period of 6 days. Data is presented as Mean ± SEM, n=6, statistical significance determined by unpaired, nonparametric, two-tailed Mann Whitney test. **p<0.01, ****p<0.0001. (**D**) Histopathology of the cross section of the ileum stained with H&E on day 6 post-wounding. Scale bars: 150μm. (**E**) Assessment of the inflammation score in mice treated with 2% brequinar compared to controls in the terminal ileum. Data is presented as Mean ± SEM, n=6, statistical significance determined by unpaired, nonparametric, two-tailed Mann Whitney test. **p<0.01. (**F**) Histopathology of the cross section of the ileum stained with H&E on day 6 post-wounding in mice treated with isotype control or anti-mouse Gr-1 antibody for a period of 7 days (100μg/mouse daily intraperitoneal injection). Scale bars: 150μm. (**G**) Assessment of the inflammation score in the ileum of mice treated with anti-mouse Gr-1 antibody and isotype controls. Data is presented as Mean ± SEM, n=7, statistical significance determined by Kruskal-Wallis test. **p<0.01 and ***p<0.001.

In mice treated with isotype antibody, the histopathology scores were elevated in brequinar-treated animals as compared to Aquaphor-treated controls (**Figure 8F and 8G**). Gr-1 antibody administration in topical 2% brequinar-treated mice reduced the inflammation in the ileum; however, it did not completely prevent the onset of inflammatory response (**Figure 8F and 8G**). While the epithelial layer and villi length were improved after Gr-1 antibody administration in 2% brequinar-treated mice, there remained significant presence of inflammatory infiltrate, loss of goblet cells and thickening of the muscularis propria (**Figure 8F**). These results indicate that while granulocyte activation contributes to the skin-gut crosstalk in this PG model, it is not the only mechanism by which skin ulcers drive the development of inflammation in the gut.

## Discussion

PG is the most damaging of the cutaneous extraintestinal manifestations of IBD and leads to repeated hospitalizations, increased morbidity (hazard ratio=1.72) and even mortality (52-57). Treatment strategies are inadequate, and traditional therapies employed to manage PG can aggravate intestinal inflammation and vice versa. Thus, the investigation of underlying disease mechanisms in PG is imperative to identify novel targets for therapy. The current lack of preclinical animal models represents a major limitation to the understanding of the pathogenic mechanisms contributing to the concomitant skin-gut inflammatory crosstalk. In this study we introduce a novel animal model displaying a severe neutrophilic dermatosis that not only mimics the skin lesions of human PG but also develops spontaneous intestinal inflammation. We further demonstrate that inhibition of pyrimidine synthesis in murine skin wounds is essential to the development of a murine PG-like condition.

Several studies have tried to characterize factors involved in PG pathogenesis (9, 58). Trauma to the skin, which almost invariably precedes the disease, apparently causes the release of danger signals and cytokines that contribute to uncontrolled skin inflammation and ulceration (59), but the exact mechanisms by which damage associated molecular patterns contribute to disease progression in PG is unknown. Presence of IL-8 and IL-36 in early PG lesions suggests that damaged keratinocytes start the inflammatory cascade followed by recruitment of neutrophils and other immune cells, which further exacerbates inflammation and subsequent skin damage (9). A recent study looking at the gene expression profile in perilesional epidermis and dermis showed that neutrophil degranulation and cytokine-cytokine receptor interactions are key upregulated pathways in PG (60). Current evidence points toward dysregulated innate and adaptive immune functions, mainly involving neutrophils and Th17-mediated inflammatory responses, as central mechanisms of PG pathogenesis (59).

Presence of dense neutrophilic infiltrate in PG ulcers creates an undermined violaceous border and loss of the epithelial layer in the region of inflammation in human disease (9). In addition to neutrophils, other immune cells like monocytes and T cells (Th1 and Th17) present in the region surrounding the ulcer are thought to be important players in disease progression (9, 61). In our mouse model, we show that inhibition of pyrimidine synthesis by PALA or brequinar drastically slows down the skin re-epithelialization process and promotes an inflammatory milieu that prevents a normal wound healing response (**Figure 1 and Figure 6**). While the ulcers in murine PG are enriched in neutrophils, F4/80+ macrophages and CD3+ T cells are also found in the perilesional regions of the skin (**Figure 2A and Figure 6D**) suggesting that additional pro-inflammatory signals produced both due to skin damage and by infiltrating immune cells contribute to the resulting PG-like phenotype. In fact, IL-1β, TNF-α, IL-6, CXCL1, MIP-1α, MIP-1β, MIP-2, GM-CSF, RANTES and IL-17A/F are also highly expressed in murine PG (**Figure 2C and 2D**). These cytokines have been shown to be key contributors in clinical PG progression and are mainly produced as a result of tissue damage in the skin (9). Thus, the inflammatory mediators detected in our model closely mimic those reported in human disease, perhaps with the exception of IL-36, whose levels were similar in diseased and control mice (data not shown) (9, 15, 38). This may be due to the timeline of disease progression in mice in comparison to humans, where IL-36 has been detected in early PG lesions. In addition to chemoattractants that facilitate the recruitment and migration of neutrophils to the ulcer site, Th17-mediated responses could also potentially contribute to enhanced neutrophil recruitment in our model.

In our model, we focused on understanding the role of neutrophils and NETs in disease pathogenesis due to their abundant presence in ulcerated skin. Multiple priming agents, like chemoattractants, cytokines, and microbial products have the capacity to activate neutrophils and induce the release of neutrophil cargo via a form of cell death called NETosis (62, 63). NETs are large extracellular structures with a web-like appearance consisting of a condensed chromatin scaffold decorated with citrullinated histones as well as externalized immunostimulatory molecules (50, 64). Studies performed in various autoimmune and autoinflammatory conditions have shown that neutrophils primed with circulating inflammatory cytokines can undergo enhanced NETosis at various sites like the skin or joints, and can perpetuate the cycle of chronic inflammation (65-67). For example, recent studies have shown that neutrophil DNA-derived NET complexes decorated with antimicrobial peptides (LL-37) lead to the activation monocytes and induction of Th17 polarizing cytokines in psoriasis (68, 69). Similarly, it has been shown that immune responses to NET-related antigens lead to the upregulation of type I interferon responses in hidradenitis suppurativa (70).

Enhanced NETosis has been observed in skin lesions of PG patients, as well as circulatory neutrophils from PAPA syndrome patients (49). Additionally, it has also been shown that a population of immature neutrophils called LDGs are responsible for enhanced NET formation in PG patients, and the pro-inflammatory cytokine IL-1β is a known inducer of NETs in patients with PG (49). Histological analysis of our PG model revealed the presence of tissue NETs (**Figure 5A and Figure 6D**). This led to the question of whether pro-inflammatory cytokines released at the ulcer site led to the production and mobilization of LDGs from the bone marrow in PG mice. We found that LDG production was increased in the bone marrow, which was reflected in the circulation of the diseased mice (**Figure 5C**). Importantly, serum CitH3 was significantly higher in PG mice compared to controls, demonstrating that neutrophils underwent enhanced NETosis in our model (**Figure 5D**). Treatment with cytokines *in vitro* showed that LDGs in both PALA as well as brequinar treated mice were primed with IL-1β to undergo enhanced NETosis suggesting that IL-1β might be the main driver of inflammation in our model (**Figure 5E and Figure 6H**). Taken together, these results suggest that pre-activated LDGs in the circulation migrate to distal tissue sites to undergo increased NETosis induced by local pro-inflammatory cues provided by the inflamed tissue environment.

In our PG model, skin wounds treated with topical pyrimidine synthesis inhibitors induced the development of spontaneous inflammation in the intestine. In topical PALA-treated mice, inflammation was mainly localized to the distal colon and the inflammatory milieu (IL-1β, TNF-α, IL-6, IL-17A/F, IP-10, and IL-33) was reflective of colitis or UC-like phenotype (**Figure 3D and Figure 4B**). There was abundant presence of neutrophils, macrophages and T cells in the inflamed colonic tissue in conjunction with increased chemoattractants such as CXCL1, MIP-2 and MCP-1 (**Figure 4A and Figure 4B**). Development of gut inflammation trailed behind inflammation in the skin suggesting the existence of a crosstalk between the two organs (**Supplementary Figure 1B**). Skin insult or wounding was required for the development of colonic inflammation in the PALA-induced model (**Figure 3F**). Intriguingly, topical brequinar-treated mice also developed intestinal inflammation; but the intestinal inflammation was localized to the ileum. While we have not fully characterized the inflammatory landscape in brequinar-treated animals, data obtained from *in vitro* NETosis experiments suggests similar mechanisms are at play in terms of neutrophil priming and exaggerated NET formation. Depletion of granulocytes in our disease model reduced intestinal damage but did not prevent the onset of inflammation indicating that likely there are other factors contributing to skin-gut crosstalk in our model (**Figure 8G**). There might be additional inflammatory cues that drive the communication between the skin and intestine that remain to be explored, such as other types of immune cells (monocytes and lymphocytes), inflammatory cytokines as well as microbial factors.

At this point, we have not explored as to why we see differences in the location of intestinal inflammation in the two pyrimidine synthesis inhibitor treatment modalities. This could be due to a variety of factors such as differences in absorption and metabolism of the inhibitors, kinetics of disease development, alterations to the intestinal microbiome, or the impact of PALA vs. brequinar on systemic immune responses. Nucleotide metabolism has been targeted using a variety of approaches to fight diseases, such as cancer, viral infections, and to alter the immune response in various disease conditions like multiple sclerosis and rheumatoid arthritis (26-30). Inhibition of different enzymes involved in *de novo* pyrimidine synthesis can alter the activated signaling intermediates involved in post translational modification of proteins, which could in turn exert immune modulatory effects (71). For example, PALA has been utilized in an *ex vivo* model of bacterial skin infection to induce enhanced antimicrobial peptide production by upregulating nucleotide binding oligomerization domain containing 2 (NOD2) signaling responses (30). Similarly, DHODH inhibitors have been utilized in viral infection models to stimulate interferon-mediated signaling mechanisms and an anti-viral response (26, 72). In addition to immunomodulatory effects, topical pyrimidine synthesis inhibition could activate various molecular mechanisms of cell death in the skin that influence wound healing kinetics and immune cell recruitment to the inflammation site (27).

Teasing apart all mechanisms of disease pathogenesis in complex diseases such as PG and IBD is a significant challenge. Nevertheless, the development of a novel murine PG model represents a significant step forward in the understanding of PG mechanisms by making available an easily inducible and reproducible model that closely mimics human PG at the phenotypic, cellular, and mediator level. In addition, our model provides researchers with a new preclinical tool that can be utilized to study inter-organ crosstalk between the skin and gut.

## Author Contributions

SJ and CM conceived, designed, and performed experiments, analyzed and interpreted data, wrote and revised the manuscript, and provided financial support for the project. AKP designed and performed experiments, helped with data acquisition as well as analysis and provided critical feedback on the revision of the manuscript. EEJ helped with experimental design, performed experiments, provided critical feedback and contributed to manuscript editing. NAR and JLS performed experiments and contributed to the revision of the manuscript. CF analyzed data and contributed to manuscript revisions. EVM, JPA and APF helped guide overall strategic planning as well as the experimental design process, data analysis and provided critical feedback on the revision of the manuscript.

## Acknowledgements

We thank Drs. Carol de la Motte, Claudio Fiocchi and Thaddeus Stappenbeck for insightful conversations and constructive criticism. We thank Nina Dvorina and Danielle Kish for their help with experimental design and troubleshooting. We are grateful to Judith Drazba and Gauravi Deshpande of the Lerner Research Institute Digital Imaging Microscopy Core, who provided assistance with confocal microscopy. This work utilized the Leica SP8 confocal microscope that was purchased with funding from the National Institutes of Health Shared Instrumentation Grant 1S10OD019972-01. We thank the Lerner Research Institute Histology Core for processing tissue sections for histology. We also thank the members of the Lerner Research Institute Flow Cytometry Core for their input on panel design and help with data acquisition. We appreciate the help from John Petrich in the Investigational Drug Pharmacy at the Cleveland Clinic to formulate the topical drugs used in this study. *N*-phosphonacetyl-L-aspartate (PALA, NSC224131) was obtained from the National Cancer Institute (NCI)/Division of Cancer Treatment and Diagnosis (DCTD)/Developmental Therapeutics Program (DTP) Open Chemical Repository (http:dtp.cancer.gov). This study was funded by a Crohn’s and Colitis Foundation Research Fellowship Award (Award# 662997; to SJ & CM), pilot funds from the Cleveland Clinic Research Program Committee (RPC Award# 551; to SJ & CM), the Pfizer Competitive IBD Grant Program (Grant# 52061263; to SJ), and the Assistant Secretary of Defense for Health Affairs, through the Peer Reviewed Medical Research Program under Award No. W81XWH-16-1-0439 / PR150299 (to C.M.). All illustrations were created in BioRender.

## Materials and Methods

For detailed reagent purchasing and use instructions, please refer to **Supplemental Table 4**.

### In vivo mouse model

Wild-type C57BL/6J (Stock No: 000664), 8-12 week old animals were purchased from Jackson Laboratories. Both male and female mice were equally distributed based on gender in the various treatment groups in this study. Single-housed mice were anesthetized and upper back fur was removed by shaving 24 hours prior to wounding. On day 0, post-anesthesia, a single full-thickness 5 mm circular wound was created down to the fascia using fine iris scissors and a 5 mm sticky tape template. The wound was created ∼0.5 cm posterior to the ears on the shaved region of the skin. The wound was positioned to prevent excessive access to grooming by mice. Topical drug formulations were compounded by the Cleveland Clinic Investigational Drug Pharmacy. Aquaphor was used a control treatment. Day 9 was selected as an endpoint because most of the control wounds treated with Aquaphor alone heal in a period 9 days. Mice were treated daily for a period of 9 days with topical 2% PALA formulated in Aquaphor or 0.5% brequinar formulated in Aquaphor. 2% topical brequinar animals were treated for a period of 6 days. To test the hypothesis whether presence of skin wounds is required for the mouse PG phenotype, 2% PALA was applied on skin without the presence of a wound for a period of 9 days. Health of the animals was monitored daily to identify signs of stress or decline. Disease activity index (DAI) assessment comprising of change in body weight, posture (normal vs. hunched), fur (normal vs. ruffled), stool consistency and evaluation of rectal prolapse (**Supplemental Table 1**) was performed in the animals every 2-3 days. Animals were photographed on even days to monitor the status of the wounds. On harvest day, tissue (skin and intestine), blood, bone marrow and stool was collected for a detailed histologic and molecular analysis using various downstream assays.

### Histopathology analysis

Skin and intestinal tissue was fixed in HistoChoice tissue fixative for 24 hours. Tissue was paraffin embedded and sectioned at the Cleveland Clinic Histology Core. Tissue sections were stained with hematoxylin & eosin (H&E) staining for histopathological analysis. Inflammation scores to evaluate tissue damage in the intestine were assessed in a blinded manner by a board certified, subspecialist gastrointestinal and hepatobiliary anatomic pathologist. Scoring parameters are listed in **Supplemental Table 2** and **Supplemental Table 3** and were adapted from Koelink *et al* (40). Immunofluorescence staining was performed to visualize immune cell infiltrates and NET components in the skin and intestine. Briefly, paraffin embedded tissue sections were deparaffinized, blocked using blocking buffer comprising of Hank’s balanced salt solution with 2% bovine serum albumin and 2% goat serum for 1 hour at room temperature in a humidifying chamber. Primary antibodies to visualize immune cell markers including neutrophils (MPO), macrophages (F4/80), T cells (CD3) and citrullinated histone H3 (CitH3) were added to the tissue sections at a 1:100 dilution in blocking buffer for an overnight incubation at 4°C. E-cadherin and K14 was used to visualize the epithelial layer in the intestine and proliferating keratinocytes in the skin, respectively. Next day, slides were washed in 1X phosphate buffered saline solution (PBS) (3 times, 5 minutes each) and tissue sections were incubated for 1 hour at room temperature using species-specific fluorophore-tagged secondary antibody (1:1000 dilution in blocking buffer). Slides were washed 3 times in 1X PBS followed by application of DAPI to visualize the nuclei. Appropriate rabbit, rat and mouse IgG controls were utilized based on the species of the primary antibody (**Supplemental Figure 5A and 5B**). Images were obtained using an inverted Leica SP8 confocal microscope using either 20X or 40X oil objective lens at 1X zoom factor. Whole tissue imaging (**Figure 2B**) was performed using the Leica DM6B microscope equipped with Leica DFC7000T camera.

### Quantification of inflammatory mediators

Multiplexed cytokine and chemokine analysis was performed using a custom kit from Meso Scale Diagnostics (MSD) selected from the mouse biomarker assay group. Prior to running the assay, skin and intestinal tissue from mice was homogenized in MSD lysis buffer with phosphatase and protease inhibitors using a bead-based homogenization technique. Protein in the tissue homogenates was quantified using the BCA protein assay and loaded at the concentration of 25μg/well. The MSD assay was performed according to manufacturer’s instructions. The custom panel included the following analytes: IL-1α, IL-1β, TNF-α, IFN-α, IFN-β, IFN-γ, IL-4, IL-5, IL-6, IL-10, Il-12p70, IL-13, IL-15, IL-17A, IL-17C, IL-17F, IL-21, IL-22, IL-23, IL-33, IP-10, GM-CSF, KC (CXCL1), MCP-1, MIP-1α, MIP-1β, MIP-2 and RANTES. Data was analyzed using the proprietary MSD immunoassay analysis software (Discovery Workbench 4.0). IL-36γ (tissue homogenates and plasma), SAA (plasma), CitH3 (plasma) and LCN2 (stool) was measured using ELISA.

### LDG isolation and flow cytometry analysis

Low-density granulocytes (LDGs) were isolated from the blood and bone marrow for quantification of LDG numbers using flow cytometry. Blood was obtained from mice post-euthanasia by performing a bilateral thoracotomy. Femur and tibia from mouse hind limbs were harvested for extraction of cells from the bone marrow using a technique described in detail by Toda *et al* (73). Blood and cells extracted from the bone marrow were diluted in 1X PBS containing 2% fetal bovine serum (FBS) and layered on the Lymphoprep gradient according to manufacturer’s instructions. LDGs in the peripheral blood mononuclear cells (PBMC) layer were removed after density gradient centrifugation (1200g at room temperature for 10 minutes). To specifically identify LDGs in the upper layer of the Lymphoprep gradient, cells were stained with live/dead fixable stain, CD45, Cd11b and Gr-1 (Ly6G/Ly6c) and quantified using flow cytometry on BD LSRFortessa™ Cell Analyzer. Flow staining buffer (1X PBS with 05% bovine serum albumin) was used for all antibody dilutions and wash steps. Appropriate single color compensation controls (One Comp eBeads) and fluorescence minus one (FMO) controls using cells were included in the flow panel. Analysis was performed on the FlowJo software (version 10.8.1). Gating was performed on live cells. The detailed strategy is depicted in **Supplemental Figure 2B**.

### In vitro NET formation assays

LDGs isolated from the bone by density gradient centrifugation were used for *in vitro* NETosis assays as described by Carmona-Rivera and Kaplan (74). Briefly, coverslips (12mm diameters) placed in 24-well plates were coated with Poly-L-Lysine (PLL) followed by 1X PBS wash (3 times, 5 minutes each wash). Cell suspensions of LDGs were prepared in phenol-red free RPMI media. 50μl droplets containing 200,000 cells were placed directly onto the center of each coverslip. Cells stimulated with phorbol 12-myristate 13-acetate (PMA) served as a positive control for NETosis (**Supplemental Figure 5D**). Cells were stimulated with recombinant IL-1β, TNF-α, IL-6 and IL-17A to assess the impact of cytokine priming in the disease model. Anti-IL-1β neutralizing antibody was added to the cell suspension to specifically identify the role of IL-1β in the LDG priming response. Incubation for NETosis assays was performed at 37°C, 5% CO_2_ (cell culture incubator) for 2 hours. After incubation, 4% paraformaldehyde (PFA) was added directly to the wells of the 24-well plate and cells were fixed overnight at 4°C. The next day, coverslips were extracted from the plate and blocked using ultrapure water with 0.1% gelatin as blocking buffer. Cells were stained with MPO and CitH3 overnight at 4°C to detect NETs, followed by 1 hour incubation with species-specific fluorophore-tagged secondary antibodies at room temperature. Washes with 1X PBS (3 washes) were performed after the primary and secondary antibody incubation. DAPI was used to visualize nuclei. Images were obtained using an inverted Leica SP8 confocal microscope using either 40X oil objective lens at 1X zoom factor. A total of 5 fields/coverslip were acquired for the assessment of NETs. Total area of NETs/field (**Figure 6E**) were quantified using the ImageJ software (version 1.53r 21).

### Granulocyte depletion

Granulocyte depletion using the anti-mouse Gr-1 antibody was performed in animals treated with topical Aquaphor and 2% brequinar. Gr-1 antibody (100μg/animal) was injected via the intraperitoneal route starting 24 hours before wounding. Animals were injected with Gr-1 antibody daily along with topical Aquaphor or 2% Brequinar treatment on mouse wounds. Animals were injected with an isotype control formulated in saline to compare the effects of granulocyte depletion in both topical Aquaphor and 2% brequinar-treated groups. Mice were monitored for signs of distress and decline in overall health. On harvest day (day 6), skin and intestinal tissue was collected for histology. Blood was collected from mice treated with Aquaphor for quantification of LDGs in circulation using flow cytometry after the daily administration of Gr-1 antibody or isotype control to confirm granulocyte depletion (**Supplemental Figure 4A**).

### Statistical analysis

Statistical analyses were performed using GraphPad Prism software (version 9.4.0). All data is presented as Mean ± SEM. The number of animals utilized for experiments is listed under each figure legend. All *in vitro* experiments were performed in triplicate. The difference between two groups was analyzed using unpaired, nonparametric, two-tailed Student’s t-Test. Mann-Whitney test was used to determine significance. For data sets with multiple treatment groups, Kruskal-Wallis test was used to determine significance. P value less than 0.05 was considered to be significant.

### Study approvals

All animal procedures were reviewed and approved prior to initiation by the Cleveland Clinic Institutional Animal Care and Use Committee.

