## Supplemental Figure 1 for "Inhibition of pyrimidine synthesis in murine skin wounds induces a pyoderma gangrenosum-like neutrophilic dermatosis accompanied by spontaneous gut inflammation"

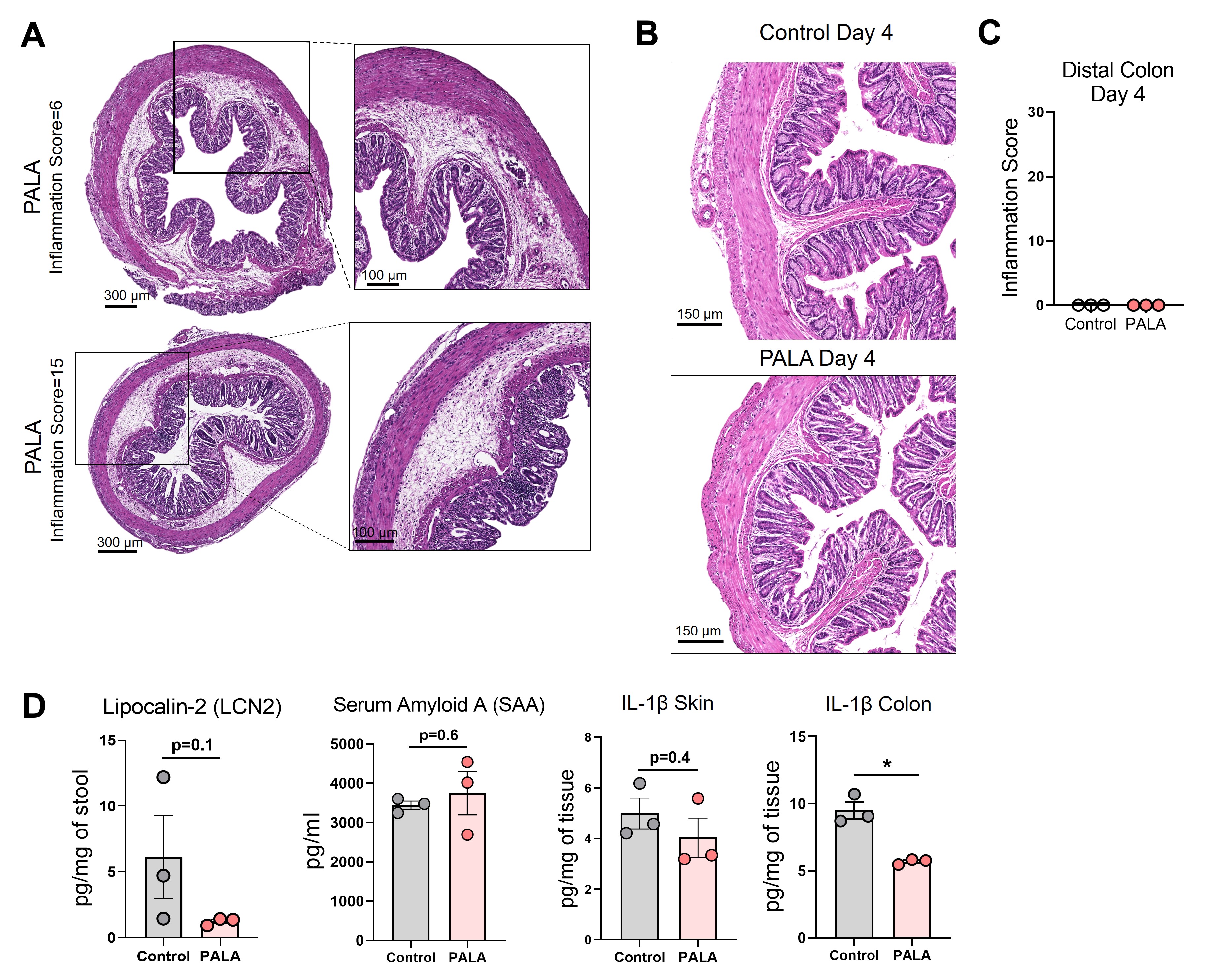

**Supplemental Figure 1.**

(**A**) Cross sections of the distal colon of mice treated with PALA with an inflammation score of 6 (top) and 15 (bottom). Scale bars: 300μm, inset: 100μm. (**B**) Histopathology of the cross section of the distal colon stained with H&E on day 4 post-wounding. Scale bars: 150μm. (**C**) Inflammation score in distal colon of mice on day 4 post-wounding. (**D**) Fecal Lcn-2, serum SAA and tissue-specific IL-1β levels in the skin as well as colon of mice in the absence of skin wound (topical PALA application only). All data is presented as Mean ± SEM, statistical significance determined by unpaired, nonparametric, two-tailed Mann Whitney test. *p<0.05. n=3.
