## Supplemental Figure 2 for "Inhibition of pyrimidine synthesis in murine skin wounds induces a pyoderma gangrenosum-like neutrophilic dermatosis accompanied by spontaneous gut inflammation"

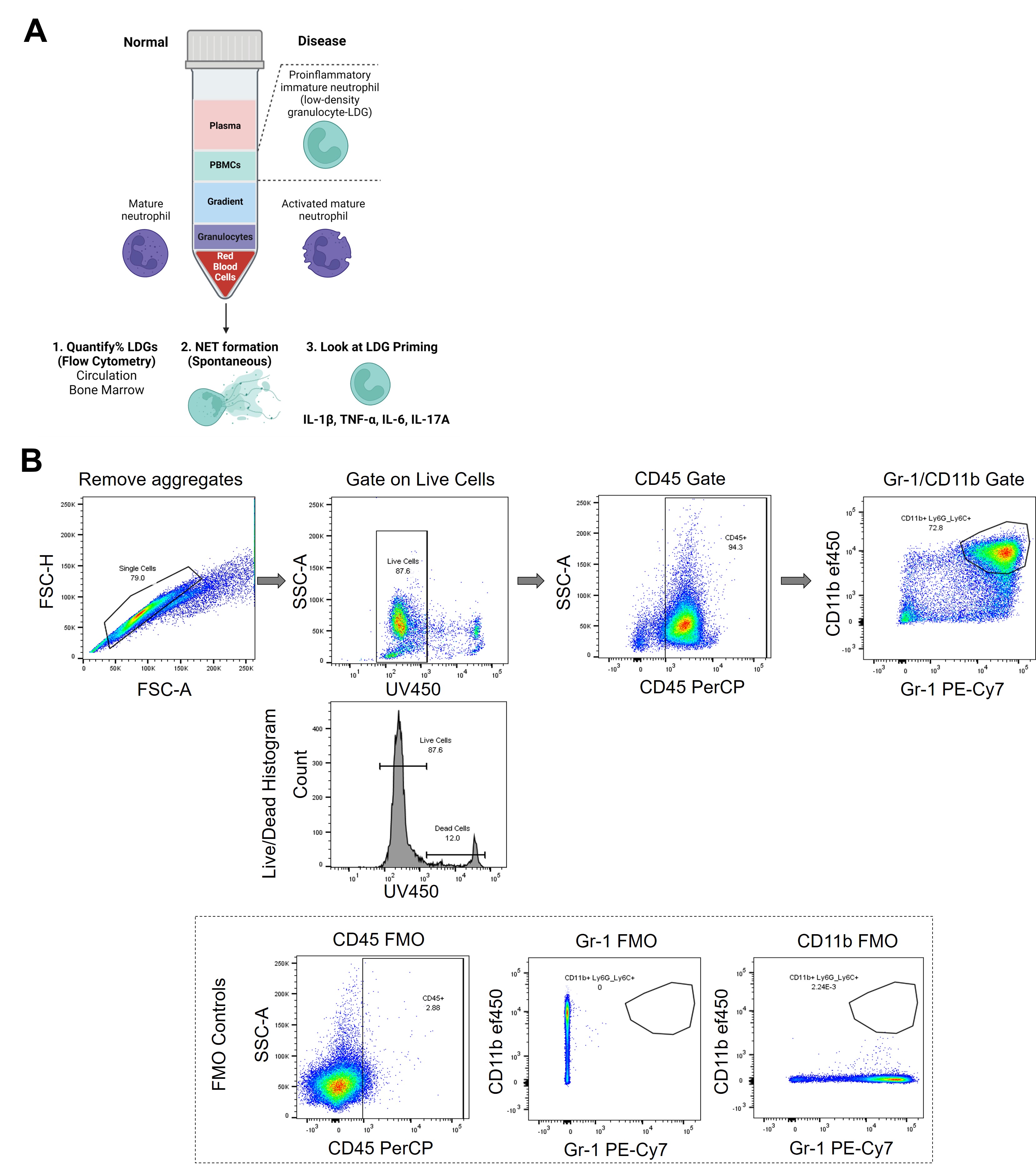

**Supplemental Figure 2.**

(**A**) Density gradient centrifugation to isolate low density granulocytes (LDGs) or immature neutrophils from blood and bone marrow for downstream applications. (**B**) Gating strategy used to quantify LDGs using flow cytometry.
