## Supplemental Figure 3 for "Inhibition of pyrimidine synthesis in murine skin wounds induces a pyoderma gangrenosum-like neutrophilic dermatosis accompanied by spontaneous gut inflammation"

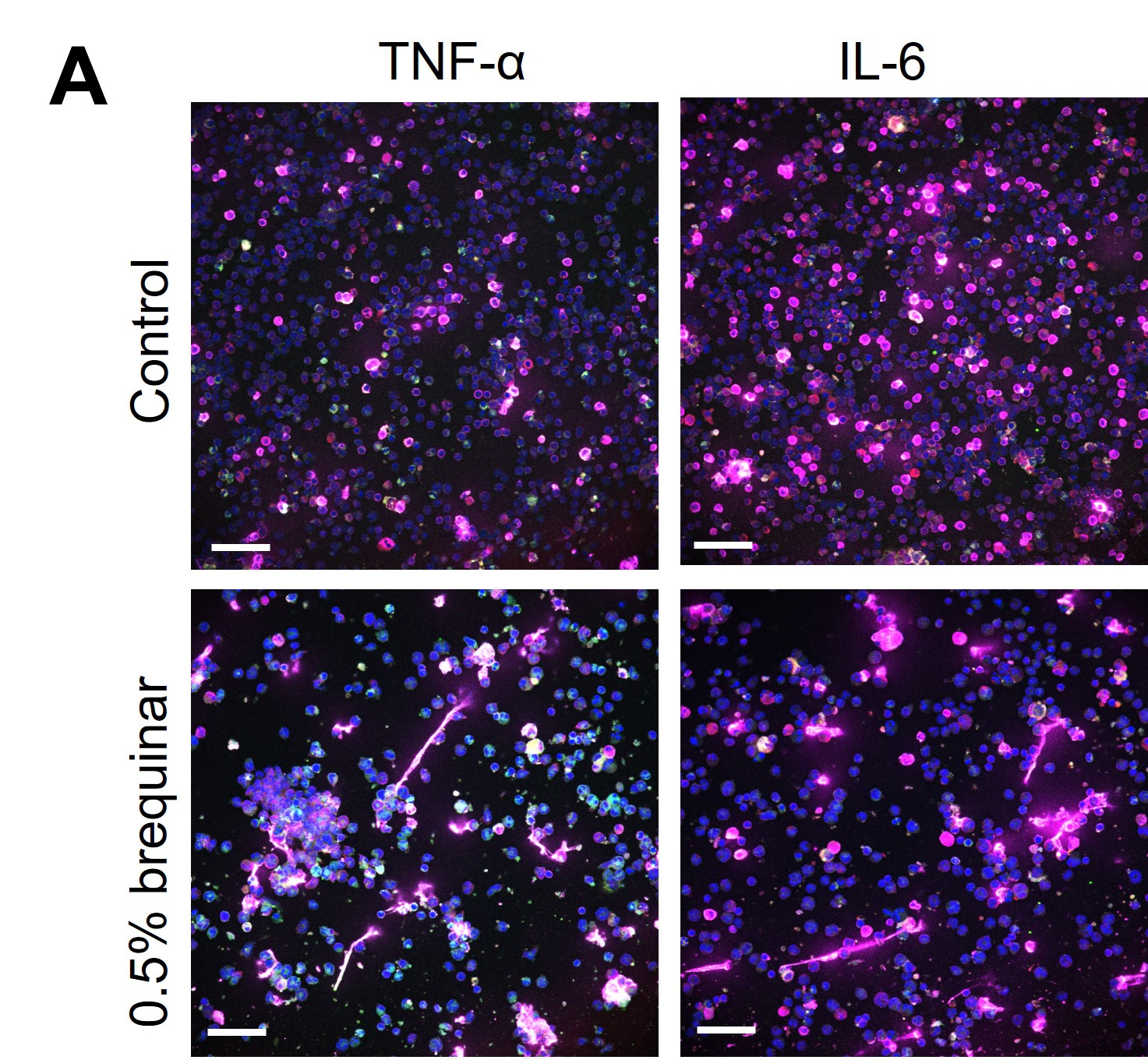

**Supplemental Figure 3.**

(**A**) Treatment of bone marrow-derived LDGs from Aquaphor-treated control mice and 0.5% brequinar-treated mice with TNF-α and IL-6 *in vitro*. NETs were stained with CitH3 (magenta), MPO (green) and DAPI (blue). Scale bars: 50μm. Images representative of n=3 mice.
