## Supplemental Figure 4 for "Inhibition of pyrimidine synthesis in murine skin wounds induces a pyoderma gangrenosum-like neutrophilic dermatosis accompanied by spontaneous gut inflammation"

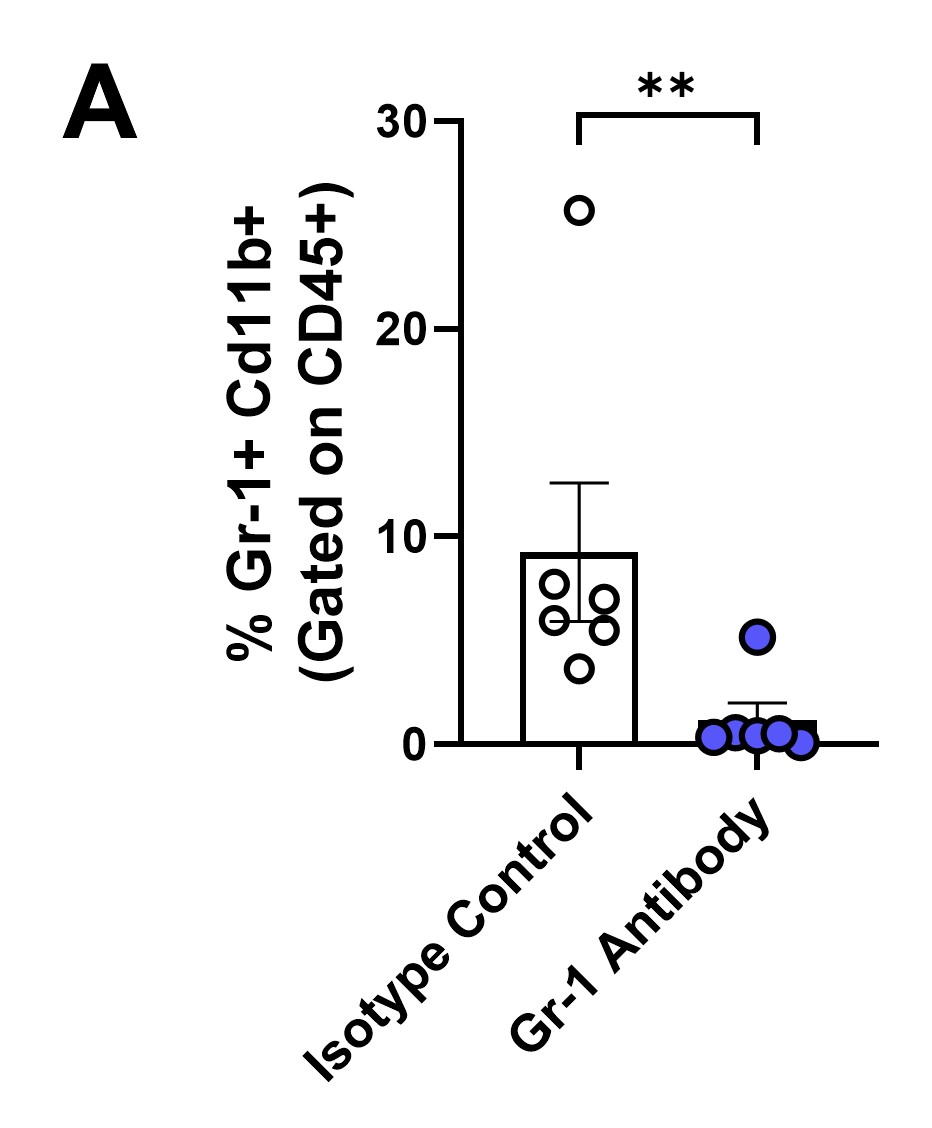

**Supplemental Figure 4.**

(**A**) Quantification of circulatory LDGs in control animals (topical Aquaphor) treated with anti-mouse Gr-1 depletion antibody and isotype controls using flow cytometry. Data is presented as Mean ± SEM, n=6, statistical significance determined by unpaired, nonparametric, two-tailed Mann Whitney test. **p<0.01.
