## Supplemental Figure 5 for "Inhibition of pyrimidine synthesis in murine skin wounds induces a pyoderma gangrenosum-like neutrophilic dermatosis accompanied by spontaneous gut inflammation"

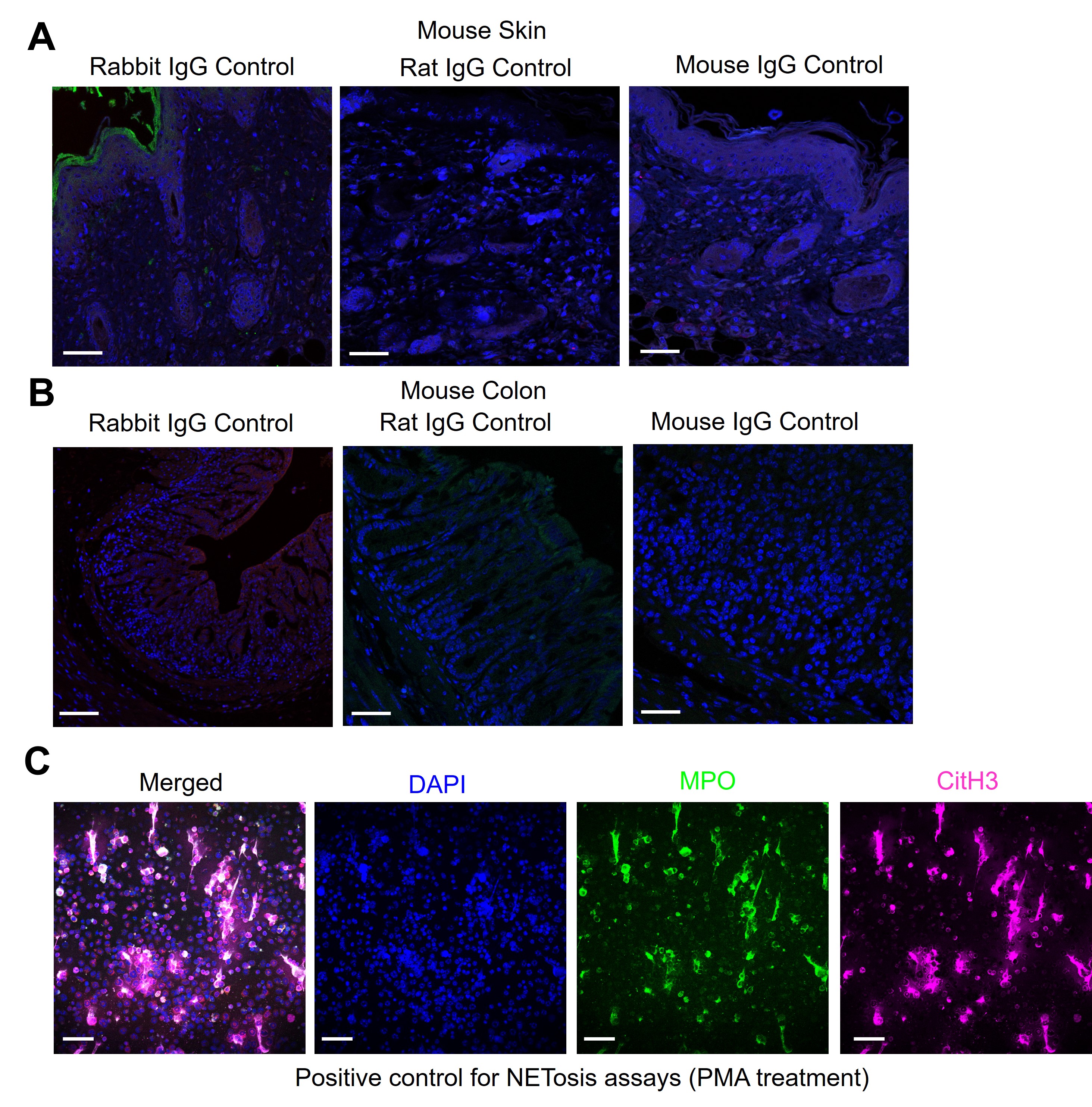

**Supplemental Figure 5**.

(**A**) IF images of IgG controls (rabbit, rat and mouse) utilized for IF staining assessment of mouse skin. (**B**) IF images of IgG controls (rabbit, rat and mouse) utilized for IF staining assessment of mouse colon. (**C**) Control LDGs treated with phorbol 12-myristate 13-acetate (PMA) as a positive control for *in vitro* NET formation assays. NETs were stained with CitH3 (magenta), MPO (green) and DAPI (blue). All scale bars are 50μm.
