## Supplementary figures and images for "Inhibition of pyrimidine synthesis in murine skin wounds induces a pyoderma gangrenosum-like neutrophilic dermatosis accompanied by spontaneous gut inflammation"

### Supplemental Table 1

**Supplemental Table 1:** Disease Activity Index (DAI) scoring parameters.

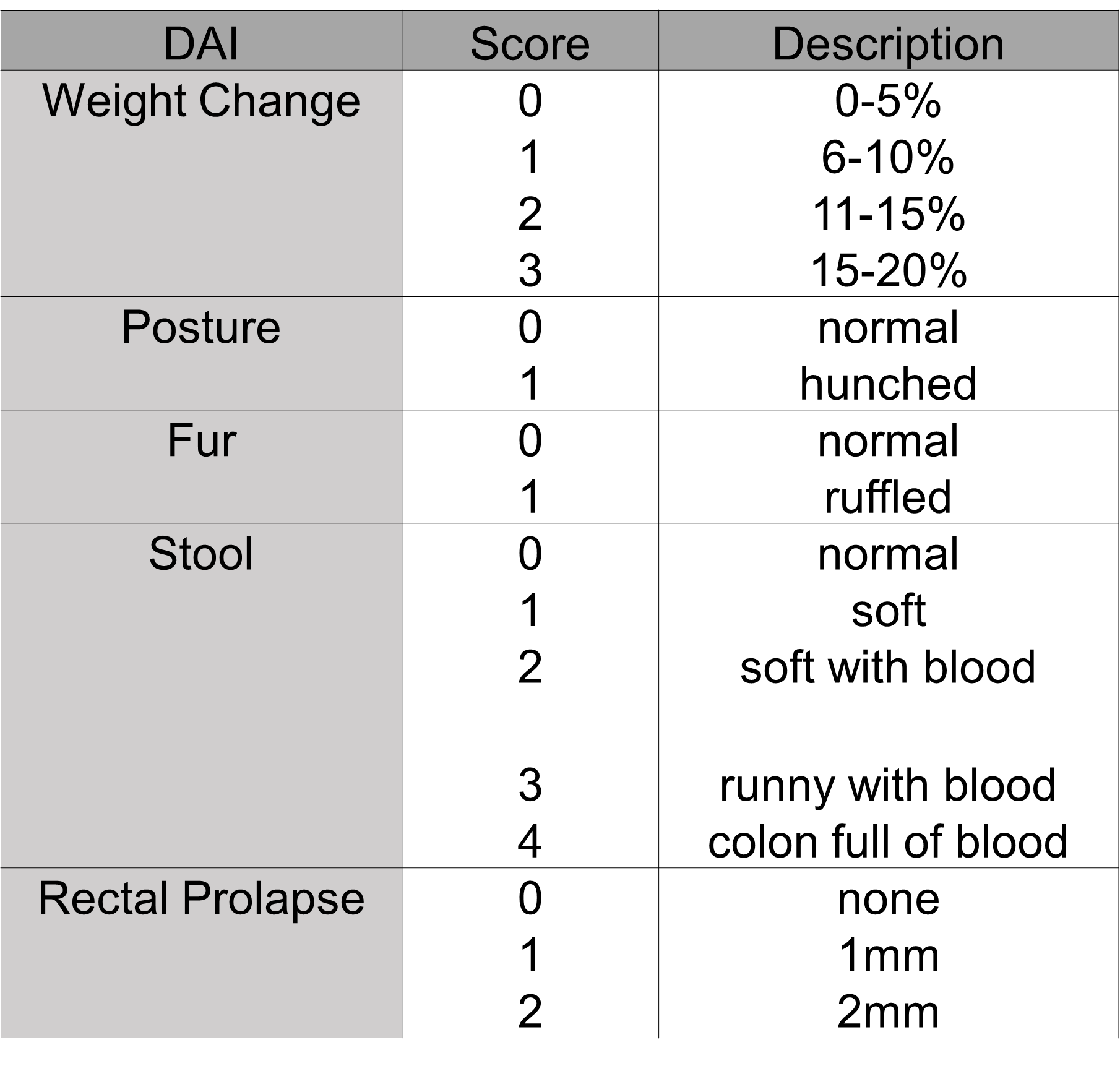

### Supplemental Table 2

**Supplemental Table 2:** Parameters used to determine inflammation score in the colon.

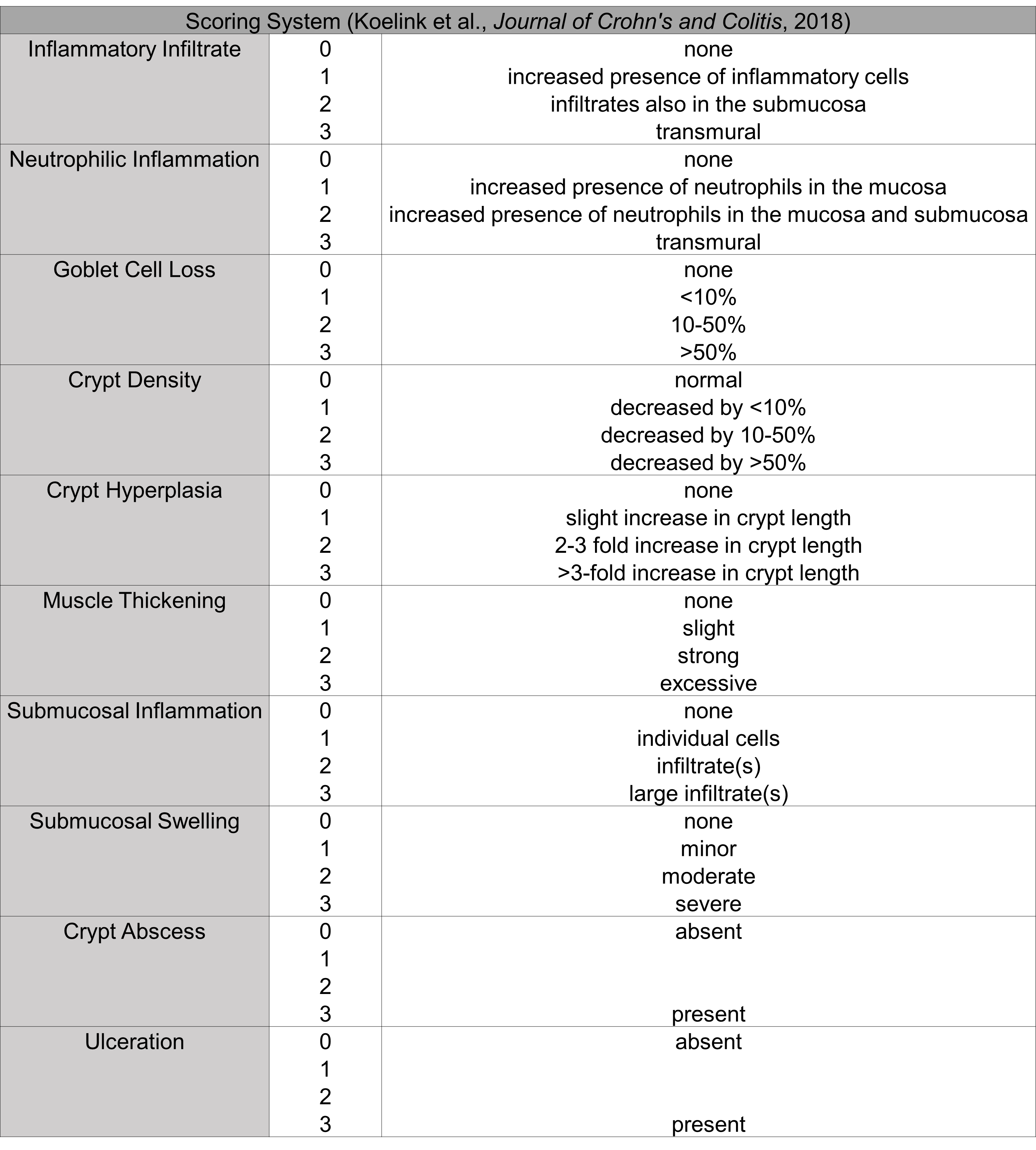

### Supplemental Table 3

**Supplemental Table 3:** Parameters used to determine inflammation score in the ileum.

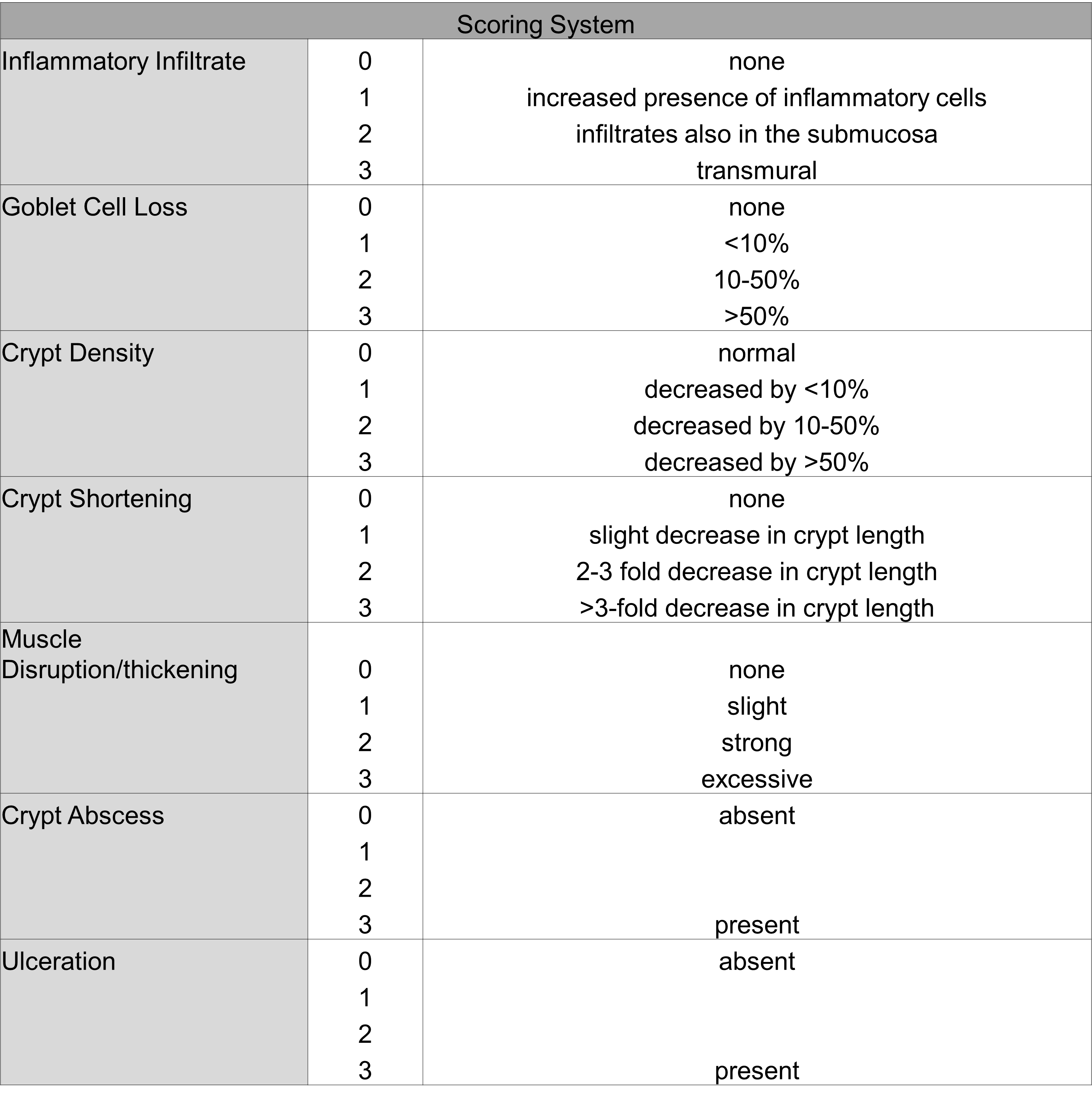

### Supplemental Table 4

**Supplemental Table 4:** Detailed reagent purchase and use instructions.

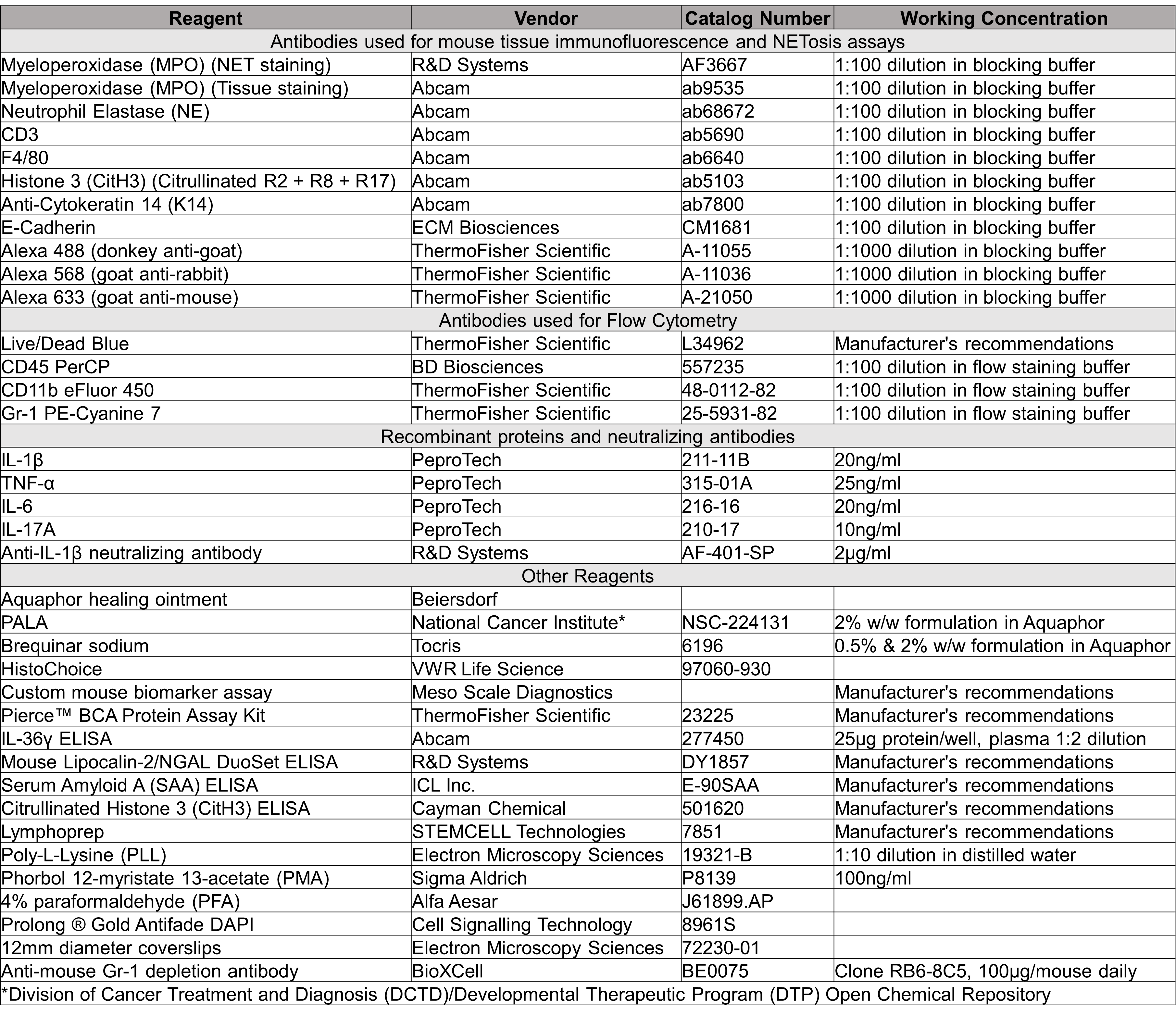
